# Droplet-based Single-cell Total RNA-seq Reveals Differential Non-Coding Expression and Splicing Patterns during Mouse Development

**DOI:** 10.1101/2021.09.15.460240

**Authors:** Fredrik Salmen, Joachim De Jonghe, Tomasz S. Kaminski, Anna Alemany, Guillermo Parada, Joe Verity-Legg, Ayaka Yanagida, Timo N. Kohler, Nicholas Battich, Floris van den Brekel, Anna L. Ellermann, Alfonso Martinez Arias, Jennifer Nichols, Martin Hemberg, Florian Hollfelder, Alexander van Oudenaarden

## Abstract

In recent years, single-cell transcriptome sequencing has revolutionized biology, allowing for the unbiased characterization of cellular subpopulations. However, most methods amplify the *termini* of polyadenylated transcripts capturing only a small fraction of the total cellular transcriptome. This precludes the detection of many long non-coding, short non-coding and non-polyadenylated protein-coding transcripts. Additionally, most workflows do not sequence the full transcript hindering the analysis of alternative splicing. We therefore developed VASA-seq to detect the total transcriptome in single cells. VASA-seq is compatible with both plate-based formats and droplet microfluidics. We applied VASA-seq to over 30,000 single cells in the developing mouse embryo during gastrulation and early organogenesis. The dynamics of the total single-cell transcriptome result in the discovery of novel cell type markers many based on non-coding RNA, an *in vivo* cell cycle analysis and an improved RNA velocity characterization. Moreover, it provides the first comprehensive analysis of alternative splicing during mammalian development.

## INTRODUCTION

Single-cell RNA sequencing (scRNA-seq) has radically transformed our understanding of cellular complexity over the last decade. Initial technologies were applied to small numbers of individual cells (Hashimshony et al., 2012; Islam et al., 2011; Ramsköld et al., 2012; Tang et al., 2009) and were subsequently adapted to droplet-based formats that can measure transcriptomes in thousands to millions of single cells (Klein et al., 2015; Macosko et al., 2015; Zheng et al., 2017). Although state of the art scRNA-seq methods are sufficiently sensitive to quantify and determine cell states with high accuracy (Grün et al., 2015; Jaitin et al., 2014; Shalek et al., 2013; Zeisel et al., 2015), most methods rely on the hybridization of barcoded oligo-dT primers to the poly(A) sequences of polyadenylated transcripts for RNA-capture and complementary DNA (cDNA) synthesis. This results in the detection of short fragments (∼ 400 – 600 bp) immediately adjacent to the poly(A)-tail of the transcript and as such remaining sequences in polyadenylated RNA molecules and the spectrum of non-polyadenylated transcripts are undetected, hindering the investigation of differential expression of non-coding RNAs, alternative splicing (AS) and alternative promoter usage (AP).

Full-length transcriptome sequencing methods (Hagemann-Jensen et al., 2020; Picelli et al., 2014) have enabled AS profiling of polyadenylated RNA species at single-cell resolution (Feng et al., 2021; Lukacsovich et al., 2019; Shalek et al., 2013), but the exact quantification of splicing events is hampered by the lack of strand and unique molecular identifier (UMI) information along the whole gene body. Furthermore, neither full-length nor whole transcriptome methods (Hayashi et al., 2018; Verboom et al., 2019) have been adapted to high-throughput droplet-based platforms, which offer at least one order-of-magnitude gain in throughput compared to plate-based methods (Svensson et al., 2018).

To overcome these challenges, we developed “Vast transcriptome Analysis of Single cells by dA-tailing” (VASA-seq) which enables the capture of both non-polyadenylated and polyadenylated transcripts across the entire transcript length. Furthermore, VASA-seq has been adapted to work with plates and droplet microfluidic formats, allowing for large-scale profiling of cell populations. To demonstrate the capabilities of our workflow, we first benchmarked VASA-seq against state-of-the-art methods using cultured cells. VASA-seq is the only technology to combine excellent sensitivity, full-length coverage of total RNA, and high-throughput in terms of number of processed cells. Next, we applied VASA-seq on a complex *in vivo* population of cells and sequenced over 30,000 single cells from mouse post-implantation embryos at developmental stages E6.5, E7.5, E8.5 and E9.5. Our resource provides the first comprehensive analysis of mammalian post-implantation development by characterizing the total transcriptome in single cells. This reveals the dynamics of alternative splicing and non-polyadenylated RNA species (*e.g.* histone, short and long non-coding transcripts) during mammalian development uncovering a layer of biological information that has been absent from recently published resources (Argelaguet et al., 2019; Cao et al., 2019; Grosswendt et al., 2020; Mittnenzweig et al., 2021; Pijuan-Sala et al., 2019). Additionally, VASA-seq exhibited higher sensitivity and detected more biotypes on average when directly compared to the dataset generated on the 10x Chromium (Pijuan-Sala et al., 2019). This led to the discovery of several cell type specific marker genes and non-polyadenylated histone gene expression patterns. The latter could be used to assign cells in S-phase, enabling the regression of cell cycle impact on transcriptomes and the characterization of cell type specific cycling kinetics.

Due to the addition of poly(A)-tails across transcript length with VASA-seq, more intronic reads were detected compared to the other methods, which allowed us to infer differentiation trajectories during development with increased confidence compared to existing resources using RNA velocity measurements (Bergen et al., 2020; La Manno et al., 2018). Finally, we used the full-length coverage to determine cell type specific splicing patterns. Both known and novel cell type specific AS patterns became apparent, and we present a comprehensive survey of AS, spanning mouse gastrulation and early organogenesis, with an emphasis on heart morphogenesis and erythropoiesis. Taken together, VASA-seq is a sensitive and scalable single-cell technology that uncovers a layer of biological information not attainable with technologies that rely on the current mRNA *termini* centric view.

## RESULTS

### VASA-seq enables detection of both non-polyadenylated and polyadenylated transcripts in single cells using plates or droplets

The first step in the VASA-seq protocol entails the fragmentation of RNA molecules from the single-cell lysate followed by end repair and poly(A)-tailing. The addition of poly(A)-tails to fragmented molecules enables cDNA synthesis from barcoded oligo-dT probes. In addition, a Unique Fragment Identifier (UFI) allows for the de-duplication of sequencing reads, enabling the accurate quantification of molecules with strand-specificity. Barcoded cDNA is amplified using *in vitro* transcription and the amplified ribosomal RNA (rRNA) is subsequently depleted. The final stages of the protocol resemble the CEL-seq workflow (Hashimshony et al., 2012) (Figure 1A). Libraries are amplified using unique dual-indexed PCR primers to enable the detection of index hopping when using the Illumina NovaSeq platform (Figure S1A).

**Figure 1.**
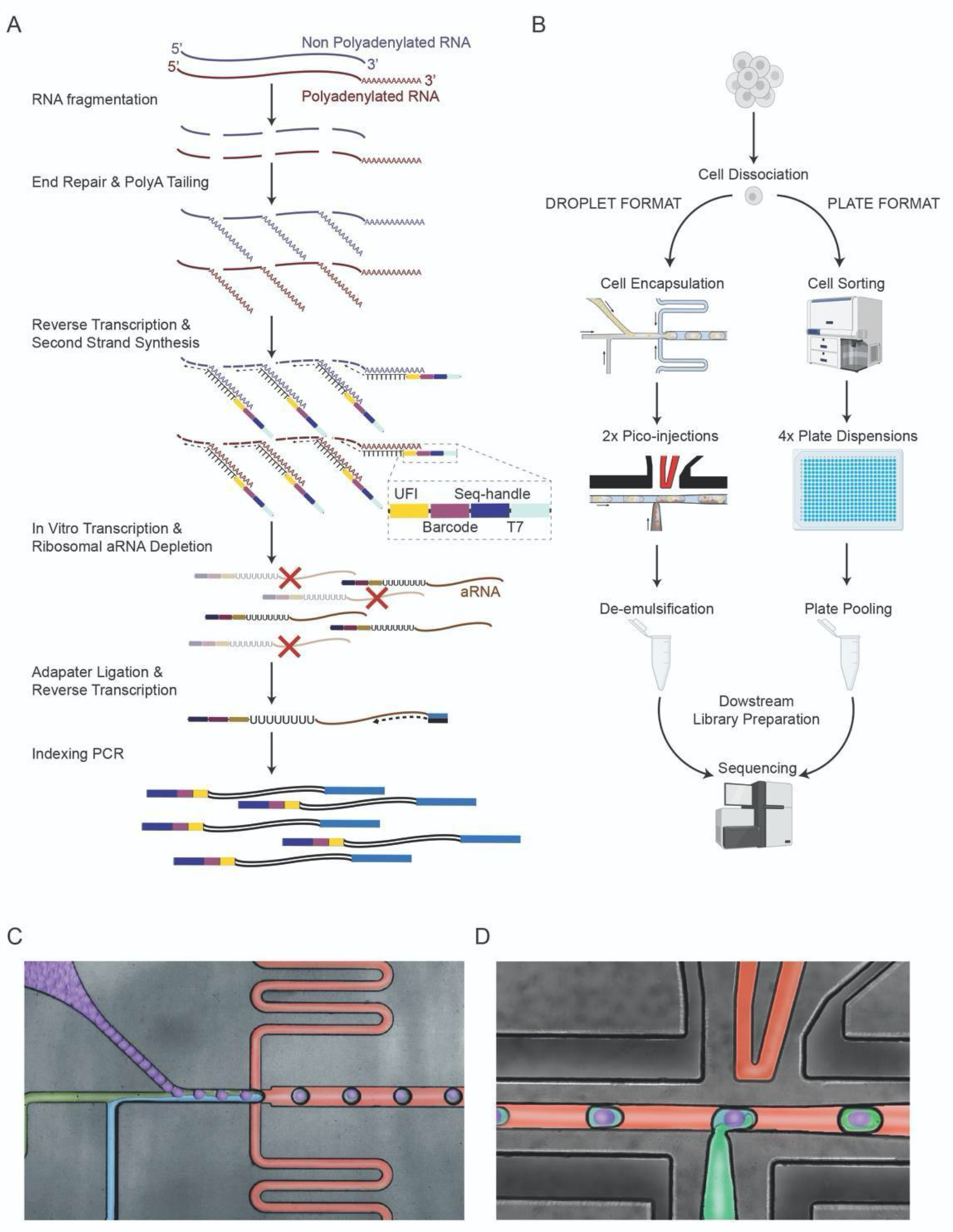
Overview of the VASA-seq workflow (A) Single cells are lysed and RNA is fragmented. Fragments are repaired and polyadenylated, followed by reverse transcription (RT) using barcoded oligo-dT primers. The cDNA is made double stranded, and amplified using *in vitro* transcription. Amplified RNA is depleted of ribosomal RNA, and libraries are finalized by ligation, RT and PCR, which leave fragments ready for sequencing. (B) The two platforms for VASA-seq, using a microfluidic device (left, VASA-drop) and a plate dispenser (right, VASA-plate). The microfluidic device allows the generation of single-cell libraries from thousands of cells, while the plate-based approach is better for rare cell types where a prior sorting is required. Each library contains the transcriptome from a large mix of cells, which are demultiplexed based on their barcode and index sequences. (C) Picture illustrating the single-cell encapsulation process using droplet microfluidics. The single cells (green) are co-encapsulated with a barcoded bead (purple), lysis and fragmentation mix (blue) and compartmentalization is achieved with the addition of fluorinated surfactant oil (red) at the flow-focusing junction. (D) Picture illustrating the pico-injection of reagents (green) to single-cell lysates (light blue/purple). The droplet surface tension is perturbed using an electric field which allows for the subsequent additions of end repair/poly(A) and RT mix.

We have adapted the VASA-seq workflow to both plate (VASA-plate) and droplet microfluidic (VASA-drop) formats (Figures 1B and S1B-G). The plate-based format is widely available and can be set up with a variety of different robots made for plate dispensation at the nanoliter scale. Plates are also beneficial when dealing with smaller numbers of rare cell types and/or when cell sorting is required. The VASA-plate workflow works by sorting cells into plates containing primers and oil (Jaitin et al., 2014; Keren-Shaul et al., 2019; Muraro et al., 2016) followed by consecutive reagent dispensing (Figure 1B). On the other hand, VASA-drop can be used for large-scale characterization of cell populations with less hands-on time and lower reagent costs. For this workflow, three microfluidic chip devices were optimized to run the reactions at high-throughput. First, a modified flow-focusing device, similar to the inDrop workflow (Klein et al., 2015), was used to co-compartmentalize cells, compressible barcoded polyacrylamide microgels and a lysis/fragmentation buffer in sub-nanoliter water-in-oil emulsions (Figures 1C and S1B). This is followed by the addition of end-repair/poly(A)-tailing and reverse transcription (RT) mixes using two consecutive steps of high-throughput reagent injections in each droplet using pico-injections (Abate et al., 2010) (Figures 1D and S1C-D). The droplets are then de-emulsified and processed for downstream library preparation.

### Barcode mixing, biotype detection, gene body coverage and sensitivity of VASA-seq

To verify that the droplet compartments remained intact throughout consecutive steps of microfluidic processing with VASA-drop, we carried out a species-mixing experiment with mouse embryonic stem cells and human HEK293T cells which showed a heterotypic doublet rate of 3.08% (Figure 2A). We then compared the VASA-seq method to the 10x Chromium droplet platform and the highly sensitive Smart-seq3 (Hagemann-Jensen et al., 2020) plate-based workflow using HEK293T cells (Figures 2A-E and S2A). Both VASA-drop and VASA-plate exhibited homogeneous coverage across the body of protein-coding genes. In contrast, 10x Chromium had the majority of its reads located near the 3’ end. Smart-seq3 had a large bias towards the 5’ end for UMI-containing reads and towards the 3’ end for the remainder of the reads (Figure 2B).

**Figure 2.**
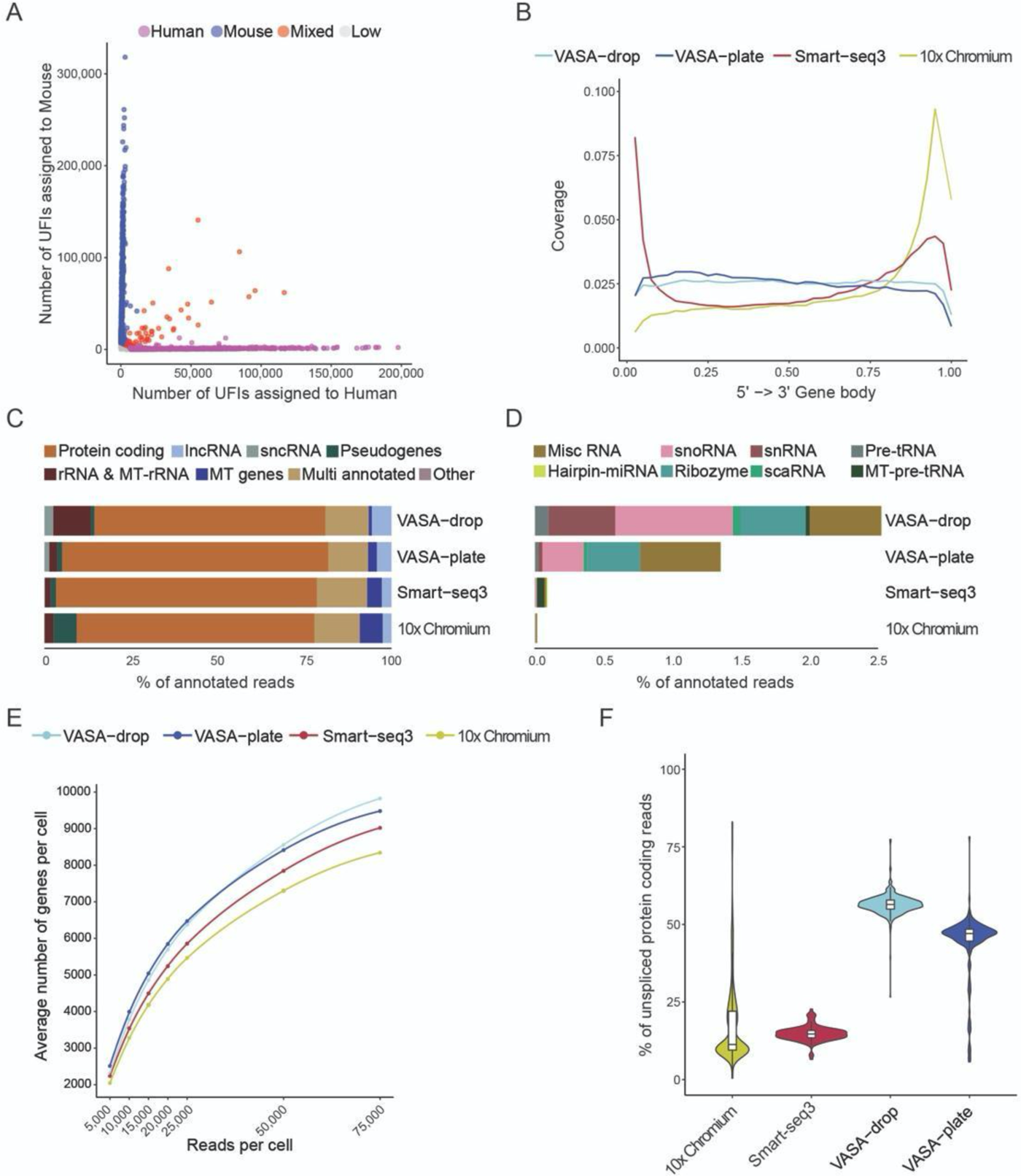
VASA-seq benchmarking against state of the art single-cell RNA-seq methods (A) Cross contamination test for VASA-drop was carried out using HEK293T (human) and mouse embryonic stem cells (mouse). Barcodes with more than 25% of detected UFIs belonging to the other species were considered doublets/mixed (red). Detected barcodes with low UFIs (<7,500) were discarded (grey). The remainder were assigned to either human (magenta) or mouse (blue). (B) Gene body coverage comparison along protein coding genes. VASA-seq showed even coverage whilst 10x and Smart-seq3 had a bias towards transcript *termini* (3’ or 5’ and 3’ respectively). (C) Biotype detection across platforms for HEK293T. VASA-seq detected approximately twice as many lncRNA genes (light blue) compared to the competitors and was the only method that captured sncRNAs at significant levels (grey). (D) sncRNA biotypes captured for HEK293T cells across methods. Only VASA-seq detected a significant number of sncRNA, withVASA-drop providing the best performance in terms of content. MiscRNA (brown), snoRNA (pink), Ribozyme(grey-green) and snRNA (red) took up the largest proportion of measured biotypes. (E) The number of detected genes in HEK293T, for each method, is plotted against the number of reads per cell across different downsampling thresholds. The saturation curves showed that VASA-seq was the most sensitive of the methods. Curvature of gene detection indicated that full complexity was not reached for the method when 75,000 reads were allocated to each cell. Only cells that were sequenced to at least 75,000 reads were used (VASA-plate: n=174, VASA-drop: n=376, Smart-seq3: n=113, 10x Chromium: n=288) (F) Percentage of unspliced reads with each method for HEK293T cells. VASA-seq detected more unspliced reads (44.1-56.5%) than the competitors (12.8-14.8%).

Protein-coding genes were the most highly detected biotype across all methods. However, VASA-plate and VASA-drop both detected about twice as many long non-coding RNA (lncRNA) as 10x Chromium and Smart-seq3 (Figure 2C). Only VASA-seq detected sncRNA, mainly miscRNA, snoRNA, ribozymes and snRNA, as well as some pre-tRNA and hairpin-miRNA (Figure 2D).

Next, the HEK293T datasets for each method were downsampled to determine gene detection sensitivity and saturation rates of each method. VASA-drop showed the highest sensitivity, followed by VASA-plate, with 9,825 ± 280 and 9,480 ± 1,252 detected genes per cell respectively (mean ± standard deviation). Both exhibited a higher gene detection rate than Smart-seq3 and 10x Chromium (9,022 ± 1,455 and 8,342 ± 1,450 genes per cell respectively) (Figure 2E). For the highest read coverage in our sequenced dataset (750,000 reads per cell, only for VASA-plate and Smart-seq3), VASA-plate and Smart-seq3 showed comparable sensitivities (15,248 ± 1,092 and 14,631 ± 988 genes per cell respectively) (Figure S2B).

Since VASA-seq detects full-length transcripts and larger amounts of unspliced RNA due to the poly(A)-tailing of RNA fragments across the transcript length, it is capable of detecting nascent transcripts at higher rates than other methodologies. To quantify this, we assigned reads that aligned to either introns or to exon-intron junctions as unspliced, whereas reads that exclusively aligned to exons were considered as spliced. VASA-seq showed the highest proportion of unspliced reads at 44.1 ± 10.1 % (VASA-plate) and 56.5 ± 3.1 % (VASA-drop), compared to Smart-seq3 (14.8 ± 2.5 %) and 10x Chromium (17.7 ± 12.8 %) (Figure 2F). Overall, VASA-seq provided unbiased full-length coverage, increased non-coding gene detection rates and the highest sensitivity in direct comparison to 10x Chromium and Smart-seq3.

### VASA-seq expands the list of cell type specific marker genes in the mouse embryo

Next, we utilized these advantages to extend and improve current atlases of mouse development. We used VASA-drop (referred to as VASA-seq in the remainder of the manuscript) to generate a single-cell total RNA-seq atlas of murine gastrulation and early organogenesis, with a total of 33,662 single cells sequenced from mouse embryonic post-implantation stages E6.5, E7.5, E8.5 and E9.5 (Figures 3A and S2C). The VASA-seq datasets from post-implantation E6.5, E7.5 and E8.5 were directly compared to a reference dataset generated using the 10x Chromium platform (Pijuan-Sala et al., 2019).

**Figure 3.**
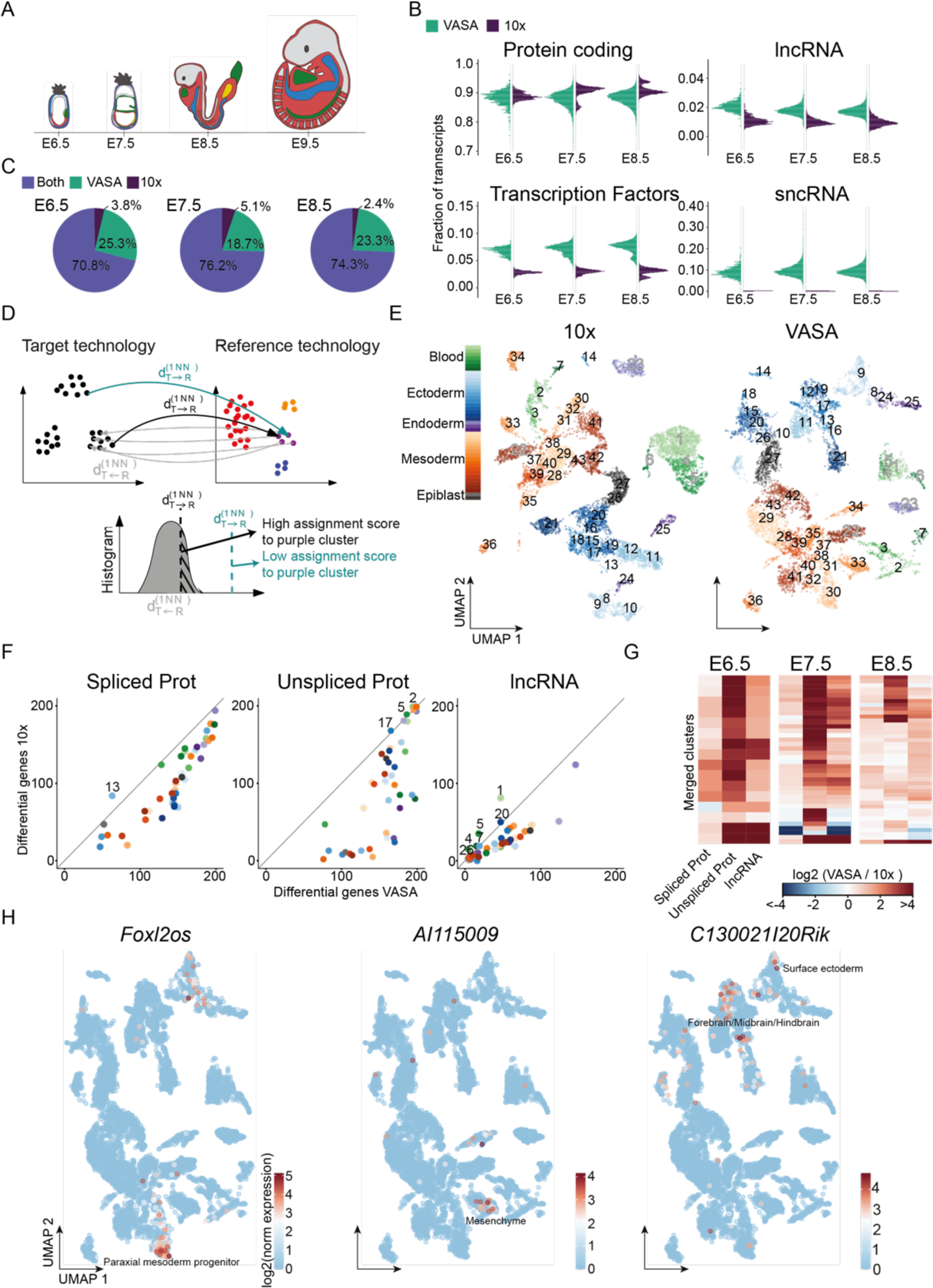
VASA-seq enables novel marker gene detection in the developing mouse embryo (A) Schematic figure of mouse embryo morphology in stages E6.5, E7.5, E8.5 and E9.5 (left to right). (B) Fraction of transcripts per biotype in VASA-seq compared to 10x for mouse embryos at each time point. The comparison includes protein coding genes (top left panel), lncRNA (top right panel), transcription factors (TF) (bottom left panel) and sncRNA (bottom right panel). (C) Percentage of genes detected in VASA-seq compared to 10x for each time point. 70.8-76.2% of the detected genes were shared between the methods, 18.7-25.3% were only detected in VASA-seq and 2.4-5.1% were only detected in 10x. (D) Strategy to transfer cluster identity from 10x or VASA (reference technology) to VASA or 10X (target technology) at the single-cell level. First, for a given cluster in the reference technology a background histogram of the distances between cells in that cluster and their corresponding first nearest-neighbor in the target technology is obtained (gray arrows and gray histogram). Next, each cell in the target technology is assigned to the cluster of its nearest-neighbor cell in the reference technology (black and green arrows) with a score equal to the area under the left curve resulting from the intersection between the cell-cell distance and the corresponding background histogram (dashed area). This procedure is then repeated for all clusters in the Reference technology. (E) UMAP of E8.5 mouse embryo cells from 10x (n=9,358) and VASA-seq (n=7,899) that were part of equivalent clusters. Clusters that are detected in both technologies are marked with numbers 1-43 and each cluster is colored according to the cell type category: green = blood, blue = ectoderm, purple = endoderm, orange = mesoderm and grey = epiblast. Grey fill in cluster label indicates extra-embryonic contribution, black fill indicates embryonic contribution. (F) Scatter plot showing the number of differentially expressed genes per cluster in VASA-seq (x-axis) vs. 10x (y-axis) for spliced protein coding (left panel), unspliced protein coding (middle panel) and lncRNA (right panel). Numbers indicate clusters where a higher number of marker genes were detected in 10x. Clusters are colored according to the cell type category: green = blood, blue = ectoderm, purple = endoderm, orange = mesoderm and grey = epiblast. (G) Heatmaps showing the ratio of differential upregulated genes (log2 fold-change above 2 and p-value below 0.01), per cluster, between VASA-seq and 10x. Columns display spliced protein coding (left panel), unspliced protein coding (middle panel) and lncRNA (right panel), whilst rows are clusters. Red color indicates when marker genes are more predominantly detected in VASA-seq, blue color indicates when higher numbers of marker genes are detected in 10x. (H) Examples of newly detected unspliced lncRNA marker genes in VASA-seq for E8.5. *Foxl2os* in paraxial mesoderm progenitors (left panel) *AI115009* in mesenchyme (middle panel) and *C130021I20Rik* in forebrain/midbrain/hindbrain and surface ectoderm (right panel).

Overall, VASA-seq detected a slightly lower fraction of protein coding transcripts, but lncRNAs and transcription factors (TF) were detected at about two- to three-fold higher levels, while sncRNAs were only captured in the VASA-seq dataset (Figure 3B). Overall, a majority of genes were identified in both methods across time points (70.8-76.2%) (Figure 3C), but 18.7-25.3% of the genes were only detected in the VASA-seq dataset whereas a much smaller fraction was observed uniquely in the 10x dataset (2.4-5.1%).

To explore whether our total scRNA-seq atlas provided more marker genes for different cell types, we identified groups of equivalent cells present in both VASA-seq and 10x and compared them through differential gene expression analysis. To achieve this, we first clustered each dataset for each timepoint independently. Second, for a given cluster and a reference technology (for example VASA-seq), a background histogram of the distances between cells in that cluster and their corresponding first nearest-neighbor in the target technology (for example 10x) was obtained. Finally, each cell in the target technology was assigned to the cluster of its nearest neighbor in the reference technology. Cells with low transfer scores were excluded and equivalent clusters with low numbers of cells in any technology were excluded from the downstream analysis (Figure 3D). Equivalent clusters between VASA-seq and 10x were defined as groups of cells with identical 10x and VASA cluster assignments (Figures 3E and S2C-D).

For E8.5 embryos, we identified 43 equivalent clusters shared between the 10x and the VASA-seq datasets, allowing for a systematic comparison and the identification of markers for respective technologies. Differential gene expression analysis was performed for each equivalent cluster for spliced/unspliced protein coding transcripts as well as for lncRNAs. Overall, VASA-seq detected a higher number of differentially upregulated genes (log2 fold-change above 2 and p-value below 0.01) for the majority of equivalent comparisons with 10x (Figures 3F-G and S2E-F). Based on the cell type annotations from Pijuan-Sala et al., 2019, examples include the detection of *Foxl2os* as a paraxial mesoderm progenitor marker, *AI115009* as a marker for mesenchyme, and *C130021I20Rik* as a specific marker for forebrain/midbrain/hindbrain and surface ectoderm (Figure 3H). Comprehensive lists of all equivalent cluster markers are presented in Table S1.

These results demonstrated that the list of marker genes obtained with VASA-seq was larger than the one obtained with 10x, especially for unspliced protein coding and lncRNA genes.

### Histone genes as *in vivo* markers for cycling cells

To further identify global gene signatures intrinsic to VASA-seq, we performed differential gene expression analysis by comparing the mean expression values for all genes across equivalent clusters and timepoints. This analysis identified a subset of genes that were significantly higher expressed in VASA-seq (22 genes, log2 fold-change > 4; p-value below 0.001), of which many were canonical histone genes, across all equivalent cluster comparisons (Figure 4A; Table S2). Consistently, most of the highly differentially expressed genes in the VASA-seq dataset are classified as non-polyadenylated (Herrmann et al., 2020) (Figure 4A, purple dots).

**Figure 4.**
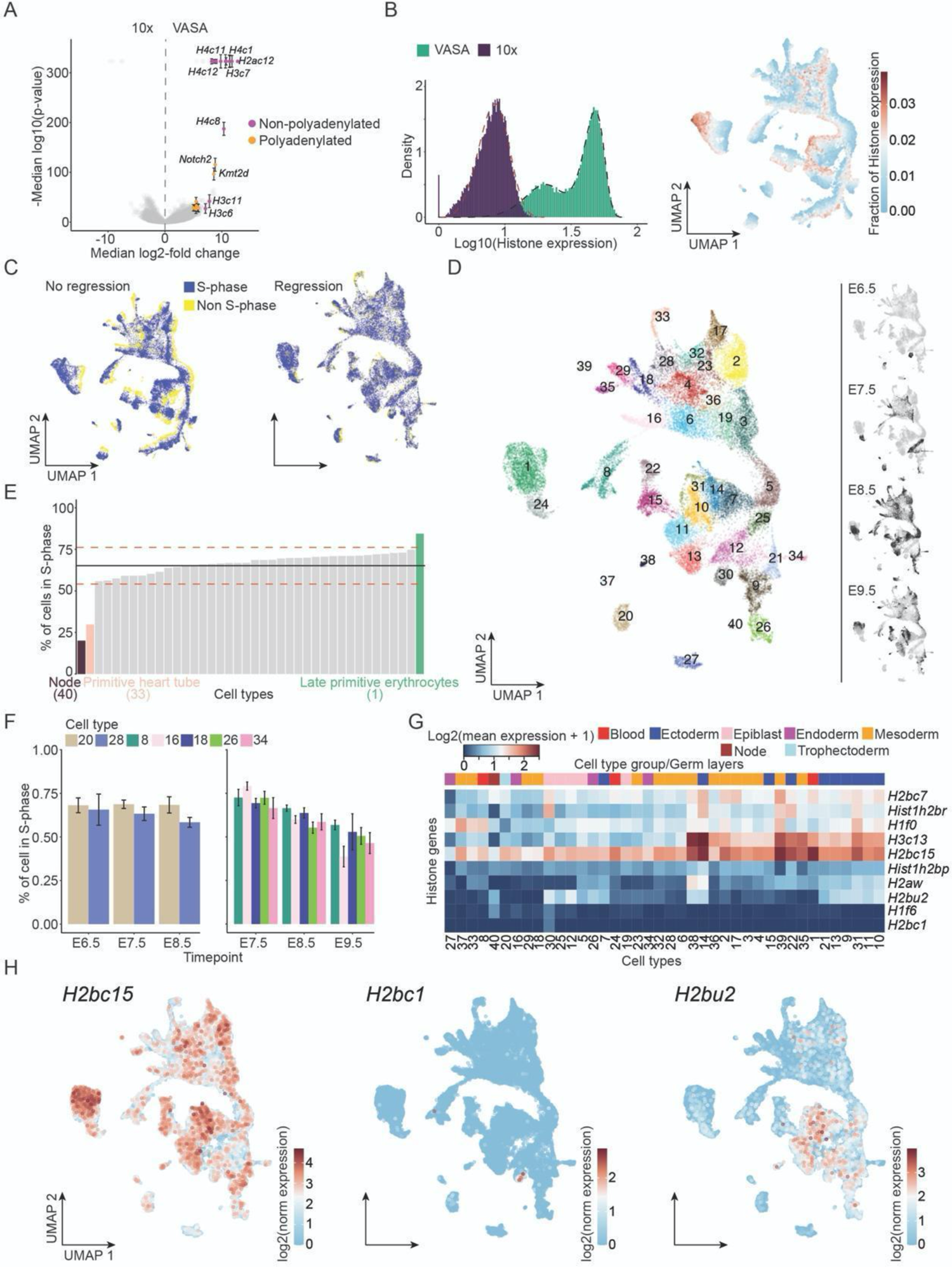
Histone gene expression robustly identifies cycling cells (A) Volcano plot showing differentially expressed genes between VASA-seq (right, positive values) and 10x (left, negative values). Genes that are always highly differentially expressed across timepoints and have a log2-fold change above 4 and a p-value below 0.001 are colored, purple color indicates non-polyadenylated and orange color indicates polyadenylated genes. Many of the differentially expressed genes enriched in the VASA-seq dataset are histone genes (B) Total fraction of histone genes in VASA-seq (red-high, blue-low), projected onto the UMAP with cells across all four time points, E6.5, E7.5, E8.5 and E9.5 (right panel) and histogram showing the distribution of histone gene expression in VASA-seq compared to 10x (left panel). Overlayed dashed black line shows a bimodal Weibull distribution and dashed red line shows a single Weibull distribution. (C) Cells are identified as cycling/S-phase (blue) and non-cycling (yellow) based on the total histone gene expression shown in Figure 4B. UMAP of the VASA-seq embryonic atlas before (left panel) and after (right panel) removal of cell cycle genes. (D) Cell-type annotated UMAP of the aggregated VASA-seq dataset after removal of cell cycle genes. Each color and number represent a cell type, called manually based on marker gene expression for each Leiden cluster. Smaller panels (right) highlight cells sampled at each timepoint (E6.5, E7.5, E8.5 and E9.5) in black. In total, 40 different cell types were identified. 1-erythropoiesis (expansive, S-phase), 2-somites, 3-paraxial mesoderm, 4-intermediate mesoderm I, 5-caudal epiblast, 6-lateral plate mesoderm/intermediate mesoderm primordium, 7-spinal cord (differentiated neurons), 8-endothelium, 9-preplacodal/placodal region, 10-rhombomeres (hindbrain), 11-forebrain/hindbrain (isthmus), 12-epiblast (E7.5), 13-forebrain, 14-spinal cord (differentiated neurons), 15-neural crest, 16-allantois, 17-cranial mesoderm, 18-lateral plate mesoderm, 19-early caudal epiblast, 20-trophectoderm, 21-dorsal surface ectoderm, 22-anterior neural crest, 23-pharyngeal arches, 24-primitive erythroid progenitors, 25-caudal epiblast (E7.5), 26-endoderm, 27-visceral endoderm, 28-first heart field, 29-myofibroblasts, 30-epiblast (E6.5), 31-spinal cord (cycling progenitors), 32-pharyngeal arches, 33-primitive heart tube, 34-outflow tract, 35-secondary heart field, 36-intermediate mesoderm I, 37-parietal endoderm, 38-pro-nephros, 39-mesodermal unknown, 40-node. (E) Percentage of cycling/S-phase cells per cell type. Average number of cycling cells is 65% (black line) ± 11% (red dashed lines) across all cell types. Late primitive erythrocytes (green) diverge from the average by having 84% of the cells in S-phase. node cells (brown) and primitive heart tube (pink) have much fewer cells in S-phase, 20% and 30% respectively. (F) Plots showing the percentage of cells in S-phase per cell type that spans over three timepoints (E6.5-E8.5, left panel, E7.5-E9.5, right panel). Trophectoderm (light brown) had an unchanged pattern while endothelium (green), allantois (pink), lateral plate mesoderm (blue), endoderm (light green), visceral endoderm (light blue) and outflow tract (dark pink) all had a decreasing fraction of cycling cells as time passes. Allantois has the biggest difference, with 38% cycling in E9.5 compared to 79% in E7.5. Error bars show values after bootstrapping. (G) Heatmap showing differentially expressed single annotated histone genes. Rows display genes, and columns display cell types. Cell type categories/germ layers can be identified by color above the heatmap. (H) Example of marker histone gene expression plotted on the UMAP, red represents high expression and blue represents low expression. *H2bc15* is highly expressed in most cell types, but absent in certain cell types, *H2bc1* is solely expressed in the early epiblast (E6.5, cell type 30), while *H2bu2* is specific to the ectoderm germ layer and epiblasts (cell types 12 & 30).

We reasoned that histone gene expression could be further used to identify cell cycle state, since the majority of canonical histone genes are strongly upregulated during the S-phase (Marzluff et al., 2008). A histogram of total histone gene expression per cell revealed a bimodal distribution for VASA-seq, in contrast to 10x (Figure 4B, left panel). We embedded all cells from the different timepoints into a single UMAP (Wagner et al., 2018) and visualized the total expression of histone genes across the dataset (Figure 4B, right panel). Cells with high histone expression were clearly segregated in the UMAP from cells with low histone expression. The bimodal distribution of histone expression in the VASA-seq datasets enabled the classification of cells as being in S-phase (high total histone expression) or non-S-phase (low total histone expression) (Figure 4C, left panel), a feature that was not detected using standard scRNA-seq cell-cycle scoring methods (Figure S3A). (Table S3). Standard cell-cycle gene markers such as Differential gene expression analysis between S-phase and non-S-phase cells was performed for either pooled or separate timepoints, which provided us with an extended list of cell-cycle genes during mouse embryonic development *Brip1, Pcna, Pola1, Top2a* or *Cdca2* were upregulated in cells annotated as S-phase (p-value < 1e-99), but the log2 fold change of expression for these genes was lower than for histone genes (below 0.7 in contrast to above 2 for histone genes).

We then regressed out cell cycle effects by removing the cell cycling genes from our dataset, and produced an improved UMAP with reduced cell cycle patterning (Figure 4C, right panel). We clustered the regressed data using the Leiden algorithm and assigned a cell type annotation to each cluster based on markers obtained through differential gene expression (Figure 4D; Table S4). Next, we investigated if certain cell types were cycling more frequently. The proportion of cells in S-phase for each cell type in the mouse embryo was 65 ± 11%. However, some cell types displayed higher proportions of cells in S-phase, such as late primitive erythrocytes (84%), while node cells and cells from the primitive heart tube showed lower proportions of cycling cells, with 20% and 30% of the cells in S-phase respectively (Figure 4E). We also explored if the percentage of cells in S-phase changed for specific cell types across the probed developmental time points. We identified seven cell types that had at least 30 cells in each of three consecutive sampled timepoints: endothelium (cell type 8), allantois (cell type 16), lateral plate mesoderm (cell type 18), trophectoderm (cell type 20), endoderm (cell type 26), visceral endoderm (cell type 27) and outflow tract (cell type 34). In this subset, only the trophectoderm showed unaltered proportions of cells in S-phase from E6.5 to E8.5 (Figure 4F, left panel). The other six cell types showed a reduction in the number of cells in S-phase across timepoints, with the allantois showing the most striking decrease from 79% to 38% between E7.5 and E9.5 (Figure 4F, right panel).

Additionally, we performed differential histone gene expression analysis between cell types (Table S5). Because histones from the same family (H1, H2a, H2b, H3 and H4) have extensive sequence similarity, not all reads could be uniquely assigned to a single histone gene. We found 10 single-annotated (Figure 4G) and 14 multi-annotated (Figure S3B) genes significantly upregulated in at least one cell type (log2 fold-change > 2; p-value below 0.01). Some histone genes showed germ layer and/or cell type specific expression. For example, *H2aw* was upregulated in the ectoderm. *H2bc15* was ubiquitously expressed in most cell types, but absent in the node (cell type 40) and the visceral endoderm (cell type 27) (Figure 4H, left panel). *H2bc1* expression was only detected in epiblast at E6.5 (cell type 30) (Figure 4H, middle panel). *H2bu2* displayed specific gene expression in the ectoderm germ layer and epiblast (cell types 12 & 30) (Figure 4H, right panel).

In conclusion, VASA-seq detected a high number of histone genes, many of which are known to be non-polyadenylated. The detection of these genes enabled robust cell-cycle classification based on their relative expression, with a marked improvement compared to current state-of-the-art methods.

### Increased intron coverage with VASA-seq allows for improved RNA velocity estimates

The large proportion of unspliced transcripts detected with VASA-seq suggested that RNA velocity profiles (La Manno et al., 2018), calculated using the ratio of unspliced-to-spliced counts for each gene, could be enhanced using this method. We therefore computed the velocities and confidence values using the scVelo package (Bergen et al., 2020) in stochastic mode for all cells across all four timepoints (E6.5-E9.5). The velocities were projected as arrows and time points as colors in the UMAP (Figure 5A). The direction of velocities clearly followed the consecutive timepoints and cell type progression in the UMAP, recapitulating previously characterized trajectories in the developing mouse embryo. To contrast the velocity profiling with 10x, we repeated the same analysis with both VASA-seq and 10x datasets using the E6.5, E7.5 and E8.5 time points. The RNA velocity vectors for VASA-seq had higher confidence metrics overall (0.84 ± 0.12) compared to 10x (0.65 ± 0.12) (Figure 5B). Next, we extracted the number of genes that contributed significantly to the RNA velocity vectors. We found that the majority of significant genes were shared between the methods (1,492). However, VASA-seq detected a large number of additional genes (1,069) that contributed to the RNA velocity vector (Figure 5C; Table S6). For the genes that were shared between both methods, we quantified the goodness of the fit (r^2^) to the gene phase diagrams with the prediction made by scVelo (Figure 5D). The stochastic model of the scVelo package fitted better to VASA-seq in terms of goodness of fit (0.74 ± 0.18) compared to the 10x data (0.38 ± 0.25). Examples of genes with a r^2^ about one standard deviation above average for both VASA and 10x are shown in Figures S3C and D. Therefore, the ability of VASA-seq to detect both spliced and unspliced transcripts with high coverage allowed for better reconstruction of the RNA velocity vectors guiding differentiation trajectories.

**Figure 5.**
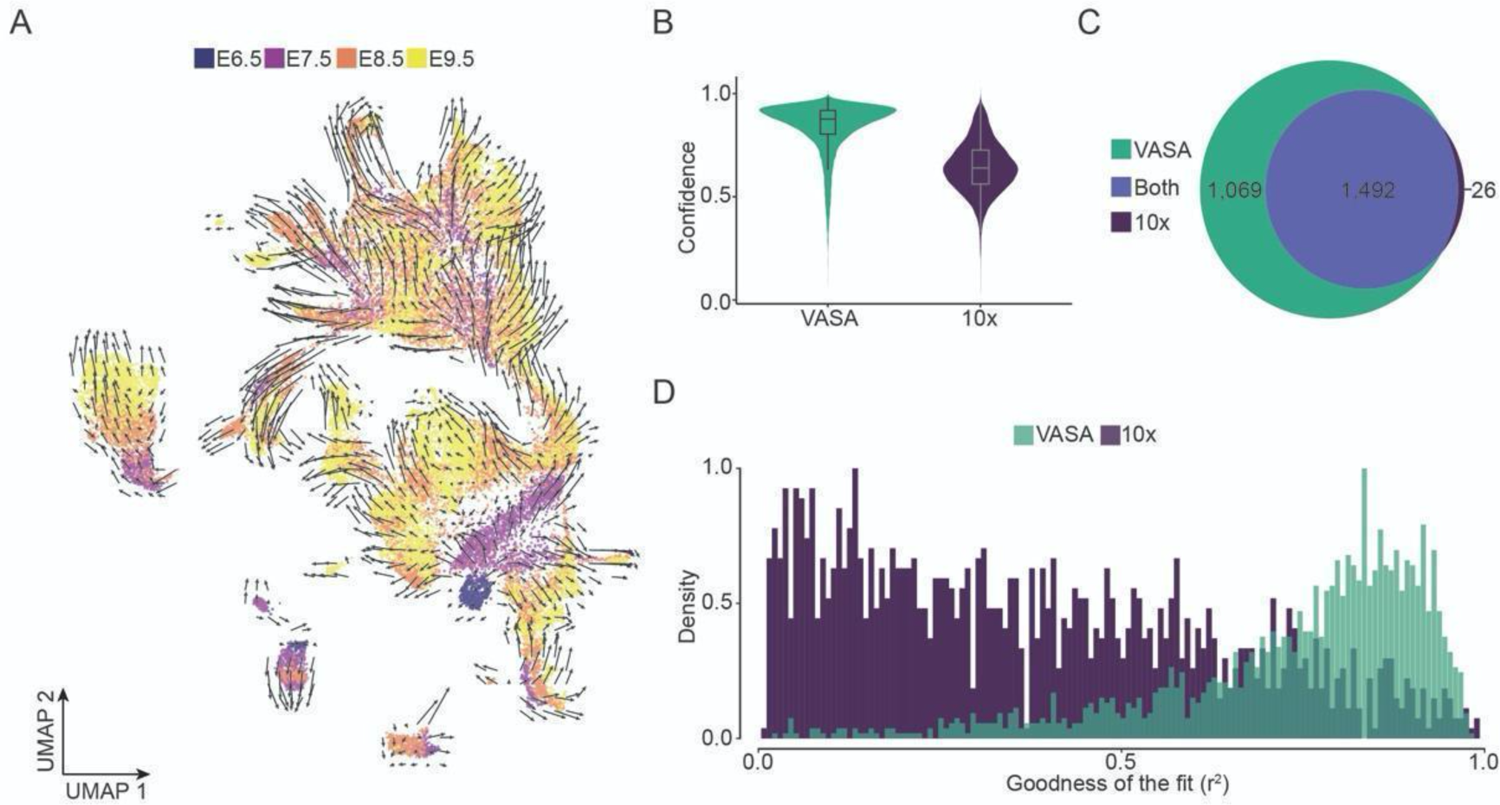
Increased intronic capture with VASA-seq improves RNA velocity measurements (A) UMAP of all four time points for VASA-seq (E6.5-E9.5). Velocity is shown as arrows and each time point as a separate color. (B) Violin plot of confidence values for VASA in green (left panel) and 10x in dark purple (right panel). Only the equivalent E6.5, E7.5 and E8.5 time points are included in the comparison. Average RNA velocity confidence was 0.84 ±0.12 (SD) for VASA-seq and 0.65 ± 0.12 (SD) for 10x. (C) Venn diagram showing the significant genes, according to the ScVelo package, for VASA and 10x. In all, we found 1,492 genes that were significant in both dataset, 1,069 that were only significant in VASA, and 26 that were only significant in 10x. (D) Histograms showing goodness of the fit (r^2^) for the 1,492 genes that were significant in both VASA and 10x. Average values were 0.74 ± 0.18 (SD) for VASA and 0.38 ± 0.25 (SD) for 10x.

### Comprehensive profiling of alternative splicing across mouse gastrulation and early organogenesis

The ability to profile full-length transcripts at scale using VASA-seq allows for the identification of alternative splicing (AS) patterns across cell types by quantifying the inclusion rates of non-overlapping exonic parts, herein referred to as “splicing nodes”. Every splicing node is associated with different types of AS, alternative transcriptional start sites or alternative polyadenylation events and their inclusion rates are calculated as percent-spliced-in (ψ) values, which is quantified by taking the ratio of reads that support the inclusion of a given splicing node (Figure 6A). To quantify AS patterns, we used Whippet, a computationally light-weight and accurate quantification method, previously integrated in computational workflows dedicated to process scRNA-seq data (Parada et al., 2021; Sterne-Weiler et al., 2018). Since splicing node coverage is limited at the single-cell level (Figure S4A), we implemented a pseudo-bulk pooling approach, developed as part of MicroExonator (Parada et al., 2021), where reads from the same cell type are pooled *in silico* before splicing node quantification. Pooling reads into pseudo-bulks from each cell-type substantially increased our ability to quantify splicing nodes (Figures S4A and B). To detect differentially included splicing nodes (DISNs) across cell types, we implemented a method developed as part of MicroExonator to detect robust AS changes across pairwise comparisons of cell-types. This method avoids the detection of spurious DISNs by arranging cells into pseudo-bulks multiple times through random sampling and integrating the results into CDF-beta values that indicate the likelihood of detecting false positive DISN events (materials and methods). Given that the total number of possible pairwise comparisons across cell clusters is prohibitively large to analyze, we aimed at detecting DISNs across cell types with closely related gene expression profiles. For this purpose, we computed the k-nearest neighbor connectivity values across cell types, and used them to generate a coarse-grained graph with PAGA (Wolf et al., 2019). The graph provided a comprehensive overview of relationships between cell clusters in terms of gene expression (Figure 6B) and enabled us to compute 72 pairwise comparisons between related cell types, from which we identified a total of 979 DISNs (Table S7). We found that 45.8% of DISNs were core exon (CE) nodes, which correspond to cassette exons involved in exon skipping, the most abundant type of AS event across vertebrates (Bradley et al., 2012).

**Figure 6.**
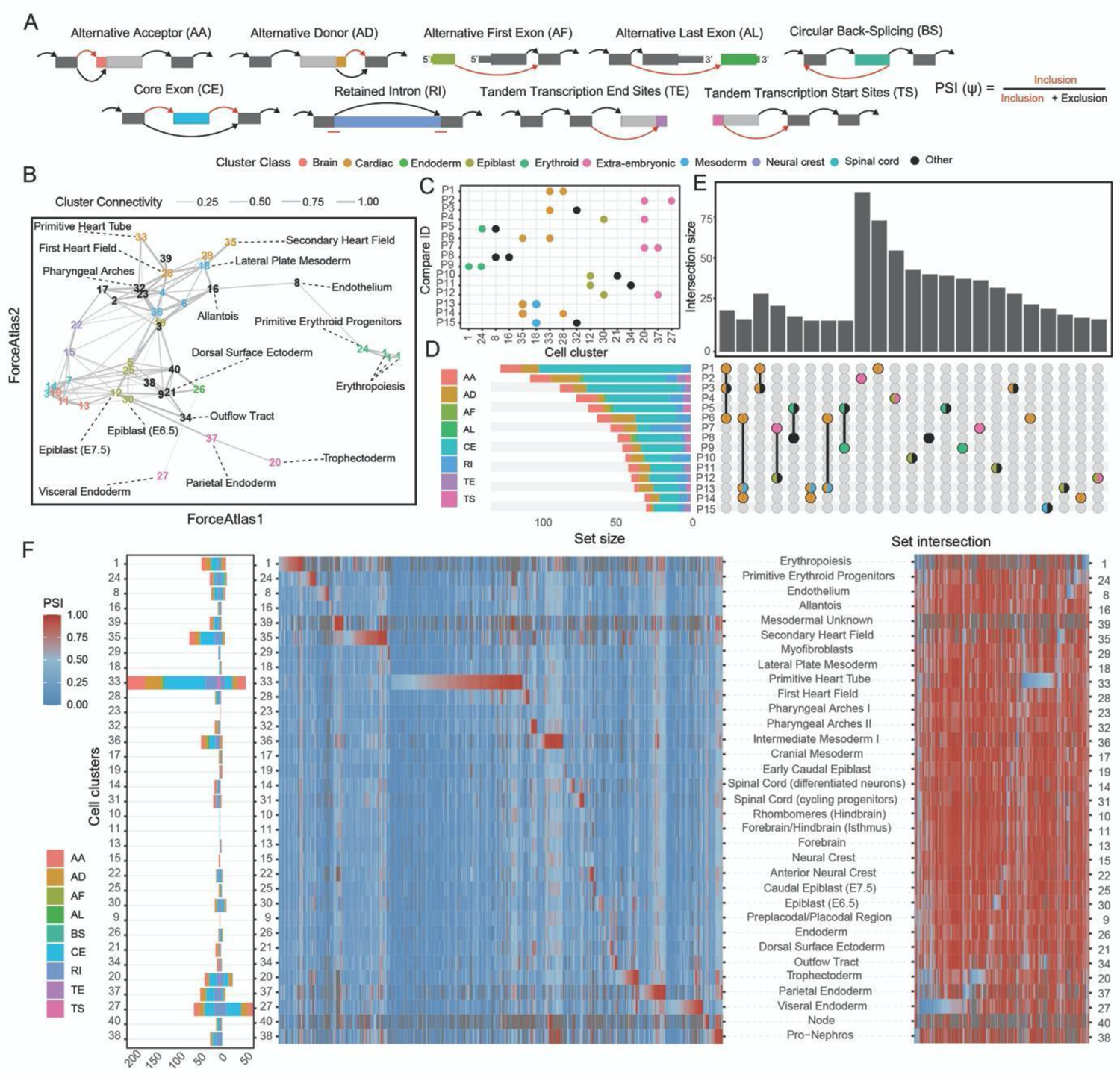
Landscape of alternative splicing events across cell types involved in gastrulation and early organogenesis (A) Schematic representation of the different types of alternative splicing nodes categorized using Whippet. Colors highlight the exonic parts that correspond to the indicated splicing nodes. Arrows indicate different possible splice junctions that can be connected to the nodes. Red arrows represent inclusion events, black arrows represent exclusion events. (B) Partition-based graph abstraction (PAGA) projecting cell types onto a force field revealing connectivity between cell-clusters. Edge width represents the connectivity values between cell clusters. Selected cell types are highlighted by color. (C) Pairwise comparisons that exhibited the highest number of differentially included splicing nodes (DISNs) are represented by a dot plot, where the position of each point on the x-axis indicates the cell clusters that were involved in each comparison (P1-P15) across the y-axis. Color codes correspond to clusters in panel B. (D) Number of DISNs found for each pairwise comparison (P1-P15). Color codes of the stacked bar plot indicate the abundance of different classes of splicing nodes that were found to be differentially included across the comparisons. (E) Upset plot showing set interactions across the group of DISNs found for each pairwise comparison. Bar plot indicates the size of each intersection, while their composition is described by the bottom panel. (F) Splice node makers identified for each cell cluster. The left panel shows the number of SNMs found for each cell cluster. The direction of the stacked bars (left or right) indicates if these markers were found with positive or negative z-score values respectively. The right panel corresponds to heatmaps displaying ψ values per cell type (blue to red scale) for different SNMs. Heatmap on the left represents inclusion SNMs and the heatmap on the right represents exclusion SNMs.

To further investigate the AS profile across cell types, we focused our analyses on the 15 pairwise comparisons that detected the highest amount of DISNs, which accounted for 67.6% of the total. These comparisons were overrepresented with cell types involved in heart morphogenesis, early gastrulation, extra-embryonic tissues and blood development, showing widespread involvement of AS in these processes (Baralle and Giudice, 2017; Conboy, 2017; Kalsotra et al., 2008; Li et al., 2020; Pimentel et al., 2013; Suckale et al., 2011). Again, CE were the most abundant type across DISNs, with the exception of comparison 7 (trophectoderm (cell type 20) vs. parietal endoderm (cell type 37)), where 40.7% of the DISNs were classified as retained intron (RI) (Figures 6C and D). Further intersection analyses between the set of DISNs revealed differential splicing nodes that were recurrently detected across different comparisons. The biggest set of common DISNs were found across comparisons that share cell clusters such as P1/P3/P6 or P6/P13/P14, which all correspond to cell types involved in heart development (Figure 6E). However, 68.7% of the identified DISNs were found exclusively across individual pairwise comparisons, suggesting a prevalence of AS events that are specific to certain differentiation transitions.

To gain further insights of global splicing patterns in relation to cell types, we identified splicing nodes with ψ values that strongly deviated from the rest of the cell types, and denoted these as splice node markers (SNMs) (Figure 6F). In total, we identified 996 SNMs (Table S8), 27.7% of which were also detected as DISNs (Figure S4C). In agreement with our previous analyses, we detected an elevated number of SNMs for cell types involved in heart development and early gastrulation. Amongst all the cell types, primitive heart tube (PHT) (cell type 33) had the most divergent splicing pattern and featured the highest number of SNMs (263), both included and excluded (Figure 6F). Moreover, we found 132 SNMs for the second heart field (cell type 35), supporting the observation of extensive AS activity during heart morphogenesis. Extraembryonic cell types that were sampled in the earlier time points mainly (E6.5 and E7.5), such as the trophectoderm (cell type 20), parietal endoderm (cell type 37) and visceral endoderm (cell type 27) also exhibited a higher-than-average proportion of SNMs (62, 56 and 132 respectively).

Taking these results together, we show that sequencing transcripts across their length with high cellular coverage using VASA-seq enabled the identification of extensive AS remodelling during mouse gastrulation and early organogenesis.

### Alternative splicing analysis of blood and heart-related tissues reveals cell type specific patterns spanning developmental trajectories

Across all cell types, the primitive heart tube had the highest number of SNMs detected and yielded the highest number of DISNs in the pairwise comparison with the first heart field (FHF) (comparison 1, Figures 6C-F). These changes occur while the heart undergoes extensive morphogenesis from E7.5 to E9.5 with the formation of the cardiac crescent consisting of the FHF and second heart field (SHF) at E7.5, that subsequently re-arranges to form the PHT at E8.0 (Später et al., 2014) (Figure 7A).

**Figure 7.**
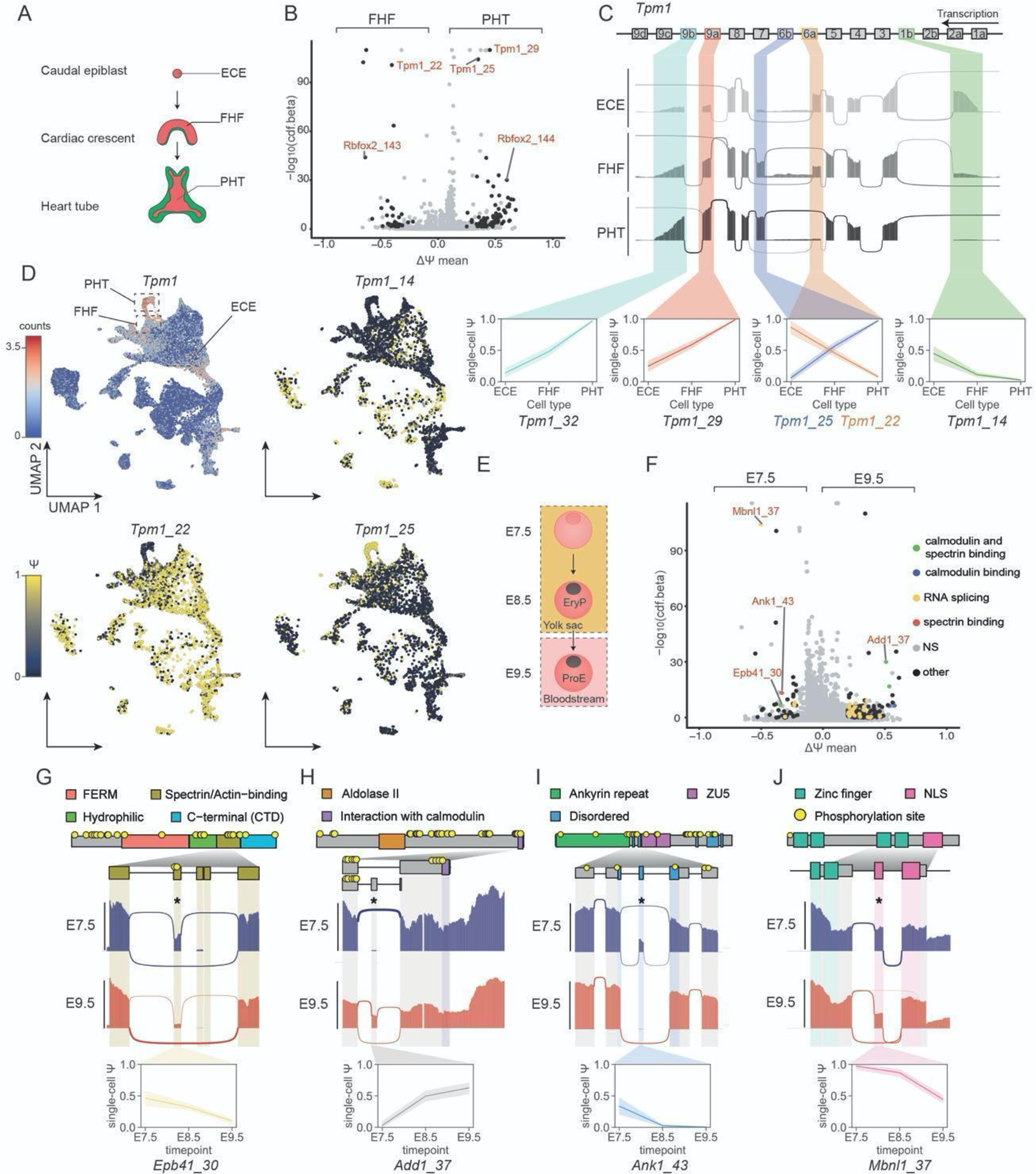
Alternative splicing patterns across heart morphogenesis and blood formation (A) Schematic of mouse heart development. (B) Volcano plot illustrating the DISNs detected between the PHT (positive ΔΨ values) and the FHF (negative ΔΨ values). (C) Sashimi plot of *Tpm1* showing a coordinated alternative splicing switch from smooth muscle to striated muscle conformation following heart development from ECE to PHT. Box annotation on top illustrates exon order for *Tpm1*. Color coding indicates the splicing node. (D) Single-cell gene expression UMAP plot for *Tpm1* (top left) and single-cell ψ projection for *Tpm1_14, Tpm1_22* and *Tpm1_25* across the global splicing analysis. Each node is color coded and highlighted in a single-cell line plot representing single-cell ψ values across all three clusters. (E) Schematic representation of murine blood development throughout the profiled timepoints. (F) Volcano plot illustrating the DISNs detected in the pairwise comparison between erythrocytes at E7.5 and E9.5. Color coding indicates proteins with calmodulin and/or spectrin binding domains or RNA splicing proteins as determined by gene ontology analysis. NS annotation stands for non-significant (grey color). (G) Sashimi, domain annotation and line plots representing the skipping of exon 16 (*Epb41_30)* between E7.5 and E9.5. (H) Sashimi, domain annotation and line plots representing the inclusion of *Add1_37* leading to a premature stop codon inclusion at E9.5 removing the C-terminus calmodulin-binding domain. (I) Sashimi, domain annotation and line plots representing the gradual exclusion of the *Ank1_43* microexon in a disordered domain. (J) Sashimi, domain annotation and line plots representing the gradual exclusion of the *Mbnl1_37* nuclear-localization signal mediating the protein’s intracellular localization across timepoints.

The detected splicing events for comparison 1 (Table S9) were coordinated with the differential expression of heart-specific RNA-binding proteins (RBP) such as *Ptbp1* (Figure S5A), that are likely orchestrating the observed AS events (Poon et al., 2012; Wei et al., 2015). In addition to changes in gene expression for RBPs, a pair of mutually exclusive exons for *Rbfox2* (*Rbfox2_143/144*, commonly referred to as B40 and M43 in the literature) were among the most significant DISNs identified in the FHF to PHT comparison (Figures 7B and S5B-C). Our results showed that B40 and M43 were preferentially included in FHF and PHT cells respectively, which is in line with previous findings depicting the inclusion of M43 over B40 in muscle-related tissues (Misra and Fisher, 2020; Nakahata and Kawamoto, 2005; Ying et al., 2017). In addition, Tropomyosin 1 (*Tpm1*) stood out with three DISNs, a CE and a pair of mutually exclusive exons, that had some of the highest confidence levels detected (*Tpm1*_29, *Tpm1*_22 and *Tpm1*_25 corresponding to exon 9a, 6a and 6b respectively). These splicing events have been shown to be part of a coordinated transition between a smooth muscle and striated muscle program, orchestrated by the antagonistic splicing regulation between PTBP1 and RBFOX2 (Cao et al., 2021). This transition was captured along a differentiation trajectory encompassing the early caudal epiblast (ECE), the FHF and PHT (Figures 7C and S5D) which also highlighted a switch for the N-(*Tpm1*_14, exon 1b) and C-(*Tpm1*_32, 9b) *termini* that has been shown to modulate the protein’s interaction with actin and troponin, hereby assisting muscle contraction (Cao et al., 2021; Hammell and Hitchcock-DeGregori, 1996). Since *Tpm1* has many cell type specific isoforms (Gooding et al., 2013), we further visualized single-cell ψ values for the aforementioned splicing nodes on the UMAP, which showed cell type specific patterning across the atlas (Figures 7D and S5E-F), underlining the potential of the VASA-seq method to capture AS at the single-cell level.

Heart morphogenesis is coordinated with the formation of blood-related cell types during the primitive blood wave (McGrath et al., 2003). At E7.25 primitive erythroids (EryP) emerge from the blood islands in the yolk sac and enter the bloodstream at E9.0 (Isern et al., 2011) (Figure 7E). The erythroid cytoskeleton then undergoes gradual rearrangements that increase their deformability when circulating in the narrow network of fetal vasculature, a change catalyzed by the adoption of the erythrocyte-specific transmembrane spectrin-actin backbone (Huang et al., 2017).

To determine if we could identify DISNs that mediate such rearrangements, we performed pairwise differential splicing analysis between erythroid cells from E7.5 (early progenitors, EryP) and E9.5 (early differentiating proerythroblasts, ProE) (Figure S5G). The analysis uncovered 210 DISNs which showed an enrichment for GO terms relating to spectrin (GO:0030507; FDR = 4.8e-03) and calmodulin (GO:0005516; FDR = 4.8e-03) binding, suggesting extensive transmembrane cytoskeletal protein rearrangements (Figure 7F; Table S10). *Epb41*, a core member of the erythrocyte cytoskeleton (Jeremy et al., 2009), showed a gradual exclusion of exon 16 (*Epb41*_30) across timepoints (Figures 7G and S4H-I). This domain contains two phosphorylation sites, directly interacts with spectrin and actin, and has been shown to be gradually included at later time points (Huang et al., 2017), suggesting a narrow exclusion window for exon 16 in primitive erythroids as they enter the bloodstream. *Add1*, which binds to ɑ- and β-spectrin and caps actin to support the membrane-bound cytoskeleton, displayed the inclusion of a premature stop codon at E9.5 (*Add1*_37) hereby excluding a C-terminal calmodulin binding domain that otherwise destabilizes its interaction with spectrin and F-actin upon calcium stimulation (Vukojevic et al., 2012) (Figures 7H and S5H-I). *Ank1,* which links the membrane to the underlying spectrin/actin filaments, had a skipped microexon (*Ank1*_43) at E9.5 that directly affected one of its intrinsically disordered regions (Figures 7I and S5H-I), which are known to be enriched in regions regulated by tissue-specific AS and predominantly contain posttranslational modification and protein-protein interaction sites (Buljan et al., 2012, 2013; Zhou et al., 2018).The identified cytoskeletal splicing rearrangements were accompanied by the detection of AS motifs in RBPs known to be involved in terminal erythropoiesis (Figure 7G, RNA splicing GO:0008380; FDR = 7.05E-06). For example, *Mbnl1* (Cheng et al., 2014), showed a skipped exon (*Mbnl1*_37) encoding for a nuclear localization signal (Figures 7J and S5H-I). Nuclear localization signal-skipping of this exon leads to its localization in the nucleus and cytoplasm rather than exclusively in the nucleus (Gates et al., 2011), likely affecting the spectrum of AS events depicted across early erythroid progenitor differentiation.

These results indicate complex AS re-arrangements that take place during mouse early organogenesis, providing functional hypotheses that can further serve as a platform for downstream validation.

## DISCUSSION

VASA-seq is a novel technology that enables the sequencing of the total transcriptome from single cells. The protocol demonstrates best-in-class capture efficiency and provides full-length coverage of coding and non-coding RNA species. In our datasets, the latter were detected at much higher levels compared to current state-of-the-art methods: 10x Chromium and Smart-seq3 (Hagemann-Jensen et al., 2020). In addition, VASA-seq does not rely on random priming which has been shown to induce sequence-specific biases in transcriptome composition (Roberts et al., 2011). Fragment quantification is also ameliorated, as it employs UMI/UFI-tagging across the whole gene body, in contrast to Smart-seq3 where only 5’ fragments are tagged; and retains strand-specificity, which improves the quantification of overlapping transcripts (Zhao et al., 2015).

The excellent performance of the method was maintained for both plate-based (VASA-plate) and droplet-based (VASA-drop) formats in our benchmarking effort, which will facilitate data integration for datasets generated from different platforms. A discrepancy arose for sncRNAs and unspliced molecules, which were detected at lower levels in VASA-plate compared to VASA-drop, possibly due to inefficient nuclei lysis in the plate experiment or different length exclusion during the DNA purification steps. On the other hand, rRNAs were not depleted as efficiently in the VASA-drop datasets, maybe because the increased barcode length for the method decreases the ability to exclude short ribosomal fragments that remain after depletion using DNA purification methods.

The throughput of the method is an order of magnitude larger than previously described total RNA-seq methodologies (Hayashi et al., 2018; Verboom et al., 2019), as the workflow is amenable to high-throughput compartmentalization and processing with droplet microfluidics. This allowed us to generate the first large-scale total-RNA-seq atlas to profile mouse gastrulation and early organogenesis. The high sensitivity and increased coverage of non-coding RNA molecules enabled us to expand the current list of cell type specific markers that will complement previous findings (Argelaguet et al., 2019; Cao et al., 2019; Grosswendt et al., 2020; Mittnenzweig et al., 2021; Pijuan-Sala et al., 2019). We further provide a detailed map of cell type specific AS events encompassing mouse development from E6.5 to E9.5, which underlined the predominance of alternative cassette exon usage throughout the time points investigated (Bradley et al., 2012; Petrillo et al., 2014; Sultan et al., 2008; Wang et al., 2008). Our resource provides the first comprehensive analysis of alternative splicing during post-implantation mammalian development.

Furthermore, VASA-seq enables the accurate estimation of cell cycle stage from direct measurements of histone content. Because most histone genes are non-polyadenylated (El Kennani et al., 2017) and since canonical histone expression is a marker for S-phase (Marzluff et al., 2008), VASA-seq outperformed previous cell cycle scoring methods based solely on polyadenylated marker expression (Scialdone et al., 2015). Accurate cell cycle determination provides especially valuable insights when the cell cycle behavior varies across developmental phases, or between different populations of cells. The workflow also enables effective removal of cell cycle effects on profiled transcriptomes, important for cell type classification and unbiased analysis (Luecken and Theis, 2019). This finding will further be useful for the benchmarking of cell cycle scoring methods, as it offers ground-truth cell cycle measurements across a breadth of different cell types and can be contrasted with the 10x chromium equivalent dataset.

Because VASA-seq captures RNA molecules across their entire length, the detected ratio of unspliced-to-spliced molecules is higher than with other methods, allowing for a more robust prediction of RNA velocity profiles. This offers a resource for further explorations that go beyond transcriptional kinetics, such as the detection of splicing dynamics across developmental trajectories. For example, our analysis of alternative splicing pointed towards extensive cytoskeletal re-modelling in maturing erythroid progenitors. However, this analysis was conducted in a supervised manner based on sequentially sampled time points, which underlines the need for unsupervised and empirical linking of velocity to AS in samples where the relationship between cell types is less understood.

The modularity afforded by the microfluidic workflow will expand the number of single-cell assays that can be performed at high-throughput. Indeed, consecutive injections of reaction mixes using droplet pico-injection modules will enable previously incompatible reactions or multi-step processes. This will benefit complex multi-omic workflows. Moreover, lower reagent costs due to smaller volumes, associated with droplet miniaturization (Salomon et al., 2019) and the lack of reliance on commercial kits in the VASA-drop workflow will enable widespread adoption of the methodology across the community.

## ACKNOWLEDGMENTS

We thank R. van der Linden for assistance during experiments. We thank Prof. Benjamin Blencowe and Prof. Patrik Ståhl, for providing valuable feedback on the manuscript. Parts of the illustrations were designed using Biorender. The work was supported by a European Research Council Advanced grant (ERC-AdG 742225-IntScOmics), Nederlandse Organisatie voor Wetenschappelijk Onderzoek (NWO) TOP award (NWO-CW 714.016.001) and Wellcome Trust (WT108438/C/15/Z). This work is part of the Oncode Institute which is partly financed by the Dutch Cancer Society. J.D.J. received scholarship support from the BBSRC, T.N.K. from AstraZeneca, A.L.E. from the Cambridge Trusts and the EU H2020 Marie Curie ITN MMBio and TSK from an EU H2020 Marie Skłodowska-Curie Actions Individual Fellowship (MSCA-IF 750772). F.H. is an H2020 ERC Advanced Investigator (69566). M.H. was supported by a core grant from the Wellcome Trust and by funding from the Evergrande Center for Immunologic Diseases. J.N. was funded by the Wellcome Trust (03151/Z/16/Z). For the purpose of Open Access, the author has applied a CC BY public copyright licence to any Author Accepted Manuscript version arising from this submission. A.Y. was funded by BBSRC (RG83885) and the mice are associated with the Wellcome Trust Strategic Grant (105031). We also thank all members of the van Oudenaarden, Hollfelder and Hemberg laboratories and Andrea Hita for scientific discussions.

## AUTHOR CONTRIBUTIONS

F.S., J.D.J., F.H. and A.v.O. designed the project; F.S. developed the VASA-seq method; F.S., J.V-L., N.B., F.v.d.B. optimized the VASA-plate workflow; J.D.J. and T.S.K. developed and optimized the VASA-drop workflow; A.Y., T.N.K., J.N. retrieved and dissociated the mouse embryos; J.D.J. and F.S performed library preparation and sequencing; A.L.E. and N.B. retrieved and dissociated the cultured cells; A.A. and F.S. developed a versatile VASA-specific pipeline and performed data analysis including benchmarking against other datasets, histone content and velocity analyses; G.P. and J.D.J. performed alternative splicing analysis and developed the associated pipelines; A.A. and A.M.A. performed cell-type annotation; M.H, F.H. and A.v.O supervised the work; F.S., J.D.J., A.A., G.P., F.H. and A.v.O. wrote the manuscript with input from all authors.

## DECLARATIONS OF INTEREST

Competing interests: F.S., A.v.O., J.D.J., T.S.K. and F.H. are inventors on patent applications submitted by Stichting Oncode Institute on behalf of Koninklijke Nederlandse Akademie Van Wetenschappen (KNAW) and The University of Cambridge (via its technology transfer office, Cambridge Enterprise).

---

**Supplementary Figure 1.**
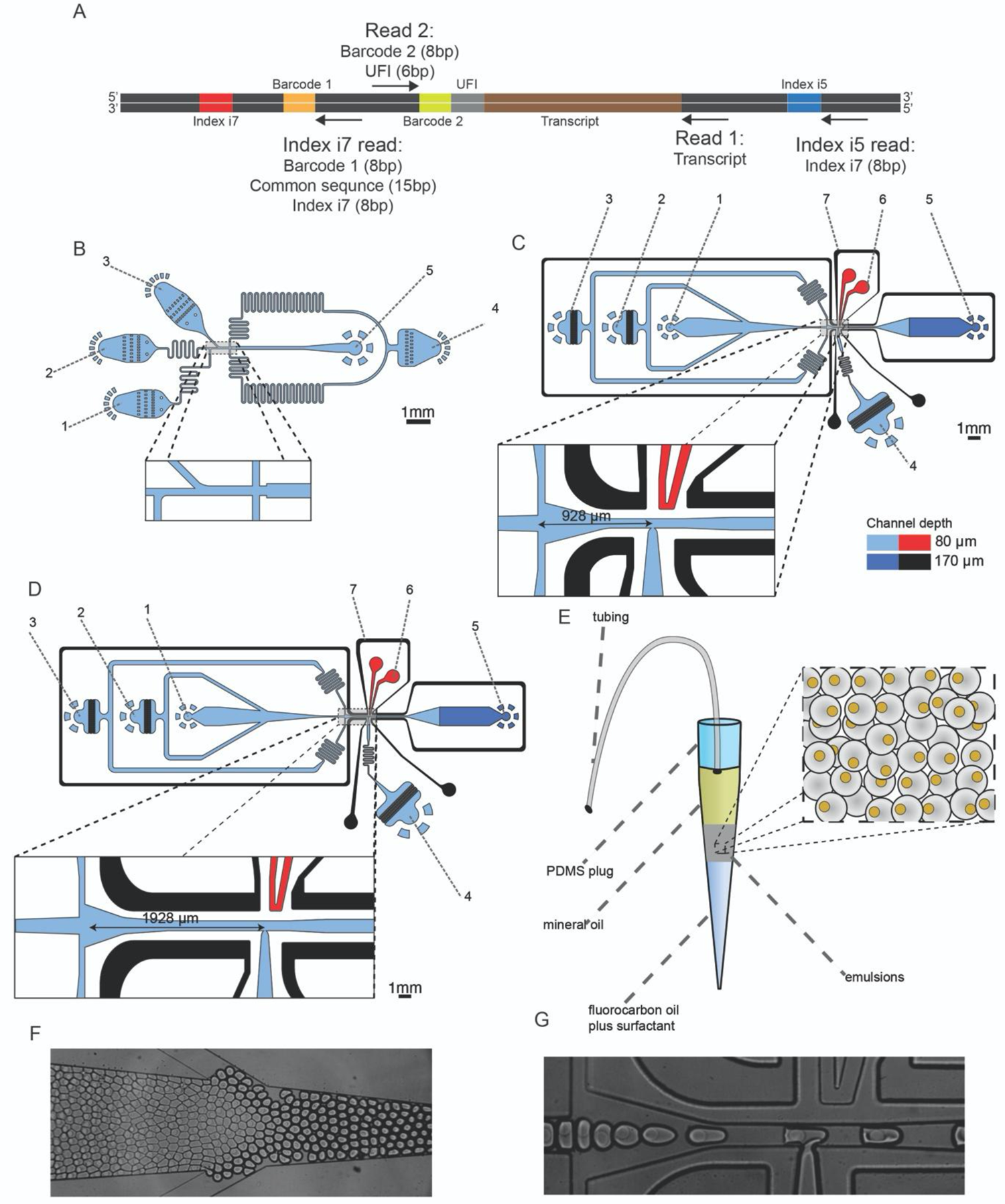
Overview of the sequencing and droplet microfluidic process A. Library barcoding design for the VASA-drop workflow. Mouse embryo libraries were sequenced with the Illumina NovaSeq platform. To avoid index hopping, a custom dual indexing strategy was used. For the index i7 read, which usually only contains barcode 1 (inDrop v3), we inserted a 8-bp second index directly after a 15bp common sequence. Only reads that had the correct combination of i5 and i7 index were further used for downstream processing. B. Design of the device used for barcoded bead and single-cell co-encapsulation. 1) input channel for the lysis and fragmentation mix, 2) input channel for the cell loading, 3) input channel for the barcoded compressible microgel loading, 4) input channel for the fluorinated oil with admixed surfactant, 5) droplet exit channel. C. Design of the first pico-injector device, to inject the repair and poly(A) polymerase. 1) input channel for droplet reinjection, 2) droplet spacing oil inlet, 3) droplet spacing oil for reinjection in the pico-injection channel, 4) inlet for the repair and poly(A) polymerase to be pico-injected, 5) droplet exit channel, 6) positive electrode (red), 7) negative (moat) electrode (black). D. Design of the second pico-injector device, to inject the RT enzyme mix. 1) input channel for droplet re-injection, 2) droplet spacing oil inlet, 3) droplet spacing oil for re-injection in the pico-injection channel, 4) inlet for the RT mix to be pico-injected, 5) droplet exit channel, 6) positive electrode (red), 7) negative (moat) electrode (black). E. Incubation chamber for droplet collection, reinjection, UV-exposure and incubation. The chamber comprises three liquid phases: 1) a top layer of mineral oil which acts as a heated lid during incubation and allows for pushing the droplets when the tubing is connected to a syringe, 2) an emulsion phase containing the single-cell lysates, 3) a fluorinated oil with admixed surfactant. Droplets can be passively collected in the chamber when the tip is inserted at the outlet of the droplet generation device and actively reinjected using a flow of mineral oil from a syringe connected though the tubing. F. Brightfield microscope images of droplet spacing before reinjection. G. Brightfield microscope images of droplet reinjection followed by pico-injection.

**Supplementary Figure 2.**
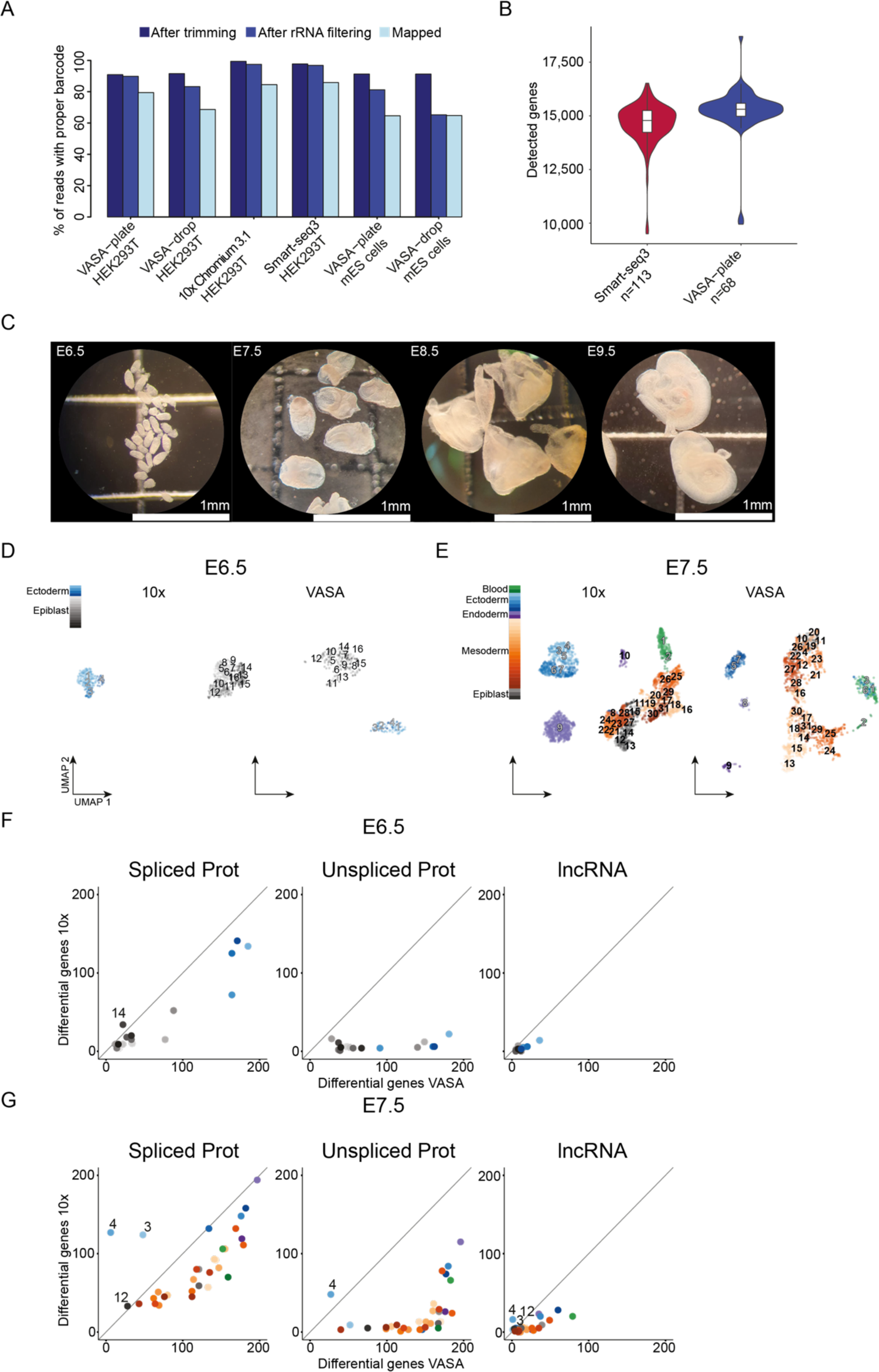
Comparison to other methods A. Percentage of sequenced reads with proper barcodes that survived trimming, rRNA filtering and mapping for each method (VASA-plate, VASA-drop, 10x Chromium and Smart-seq3) and cell type (mouse embryonic stem cells or human HEK2993T). B. Number of detected genes per cell for Smart-seq3 (red) and VASA-plate (blue) when sequenced at a depth of approximately 750,000 reads per cell. C. Brightfield microscope images of the embryos collected before dissociation. Two collections were performed for E6.5 (39 embryos total), whereas single collections were performed for E7.5 (8 embryos total), E8.5 (7 embryos total) and E9.5 (6 embryos total). D. UMAP of E6.5 mouse embryo cells from 10x (n=640) and VASA-seq (n=298) that were part of equivalent clusters. Clusters that are detected in both technologies are marked with numbers 1-16 and each cluster is colored according to the cell type category: blue = ectoderm and grey = epiblast. Grey fill in cluster label indicates extra-embryonic contribution, black fill indicates embryonic contribution. E. UMAP of E7.5 mouse embryo cells from 10x (n=3,319) and VASA-seq (n=1,892) that were part of equivalent clusters. Clusters that are detected in both technologies are marked with numbers 1-38 and each cluster is colored according to the cell type category: green = blood, blue = ectoderm, purple = endoderm, orange = mesoderm and grey = epiblast. Grey fill in cluster label indicates extra-embryonic contribution, black fill indicates embryonic contribution. F. Scatter plot showing the number of differentially expressed genes per cluster at E6.5 in VASA-seq (x-axis) vs. 10x (y-axis) for spliced protein coding (left panel), unspliced protein coding (middle panel) and lncRNA (right panel) counts. Numbers indicate clusters where a higher number of marker genes were detected in 10x. Clusters are colored according to the cell type category: blue = ectoderm and grey = epiblast. G. Scatter plot showing the number of differentially expressed genes per cluster at E7.5 in VASA-seq (x-axis) vs. 10x (y-axis) for spliced protein coding (left panel), unspliced protein coding (middle panel) and lncRNA (right panel). Numbers indicate clusters where a higher number of marker genes were detected in 10x. Clusters are colored according to the cell type category: green = blood, blue = ectoderm, purple = endoderm, orange =mesoderm and grey = epiblast.

**Supplementary Figure 3.**
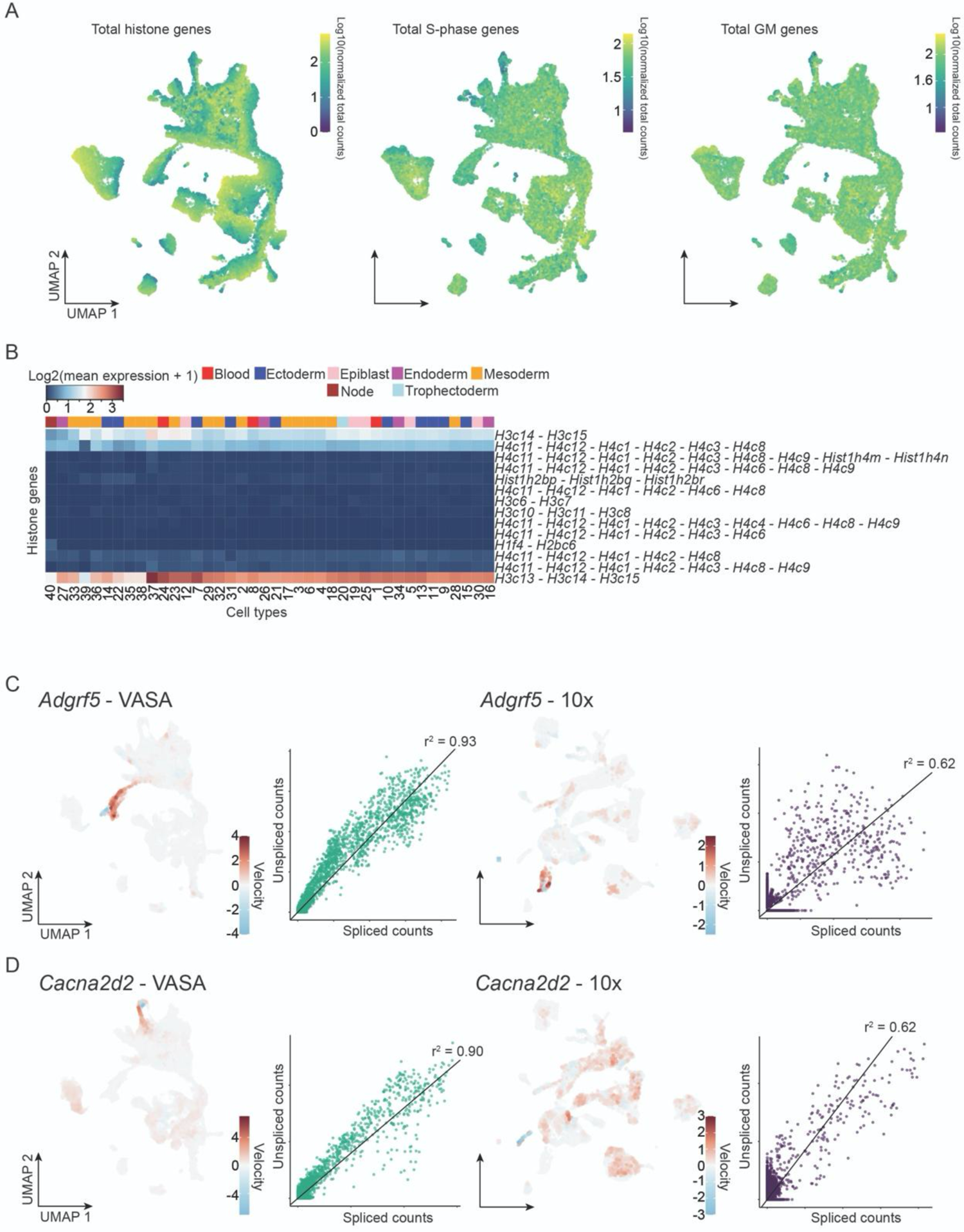
Histone expression and RNA velocity A. UMAPs showing the log10 total counts for histone genes (left panel), S-phase genes (middle panel) and GM genes (right panel). Only cell cycle scoring using solely histone genes shows a clear cell cycle segregation in VASA-seq. B. Heatmap showing differentially expressed multi annotated histone genes. Rows display genes, and columns display cell types. Cell type categories/germ layers are colored above the heatmap. C. Velocity of *Adgrf5* shown on the UMAPs for VASA-seq (left panel) and 10x (second from right panel), highlighting high velocity for the gene in the yolk sac. Phase diagrams of spliced vs. unspliced counts, with fits shown for VASA (second from left panel) and 10x (right panel). Goodness of the fit are values approximately one SD above average for each method in order to show genes that are good in both datasets. D. Velocity of *Cacna2d2* shown on the UMAPs for VASA-seq (left panel) and 10x (second from right panel), highlighting high velocity for the gene in the primitive heart tube, which was mainly observable with VASA-seq. Phase diagrams of spliced vs. unspliced counts, with fits shown for VASA-seq (second from left panel) and 10x (right panel). Goodness of the fit are values approximately one SD above average for each method in order to show genes that are good in both datasets.

**Supplementary Figure 4.**
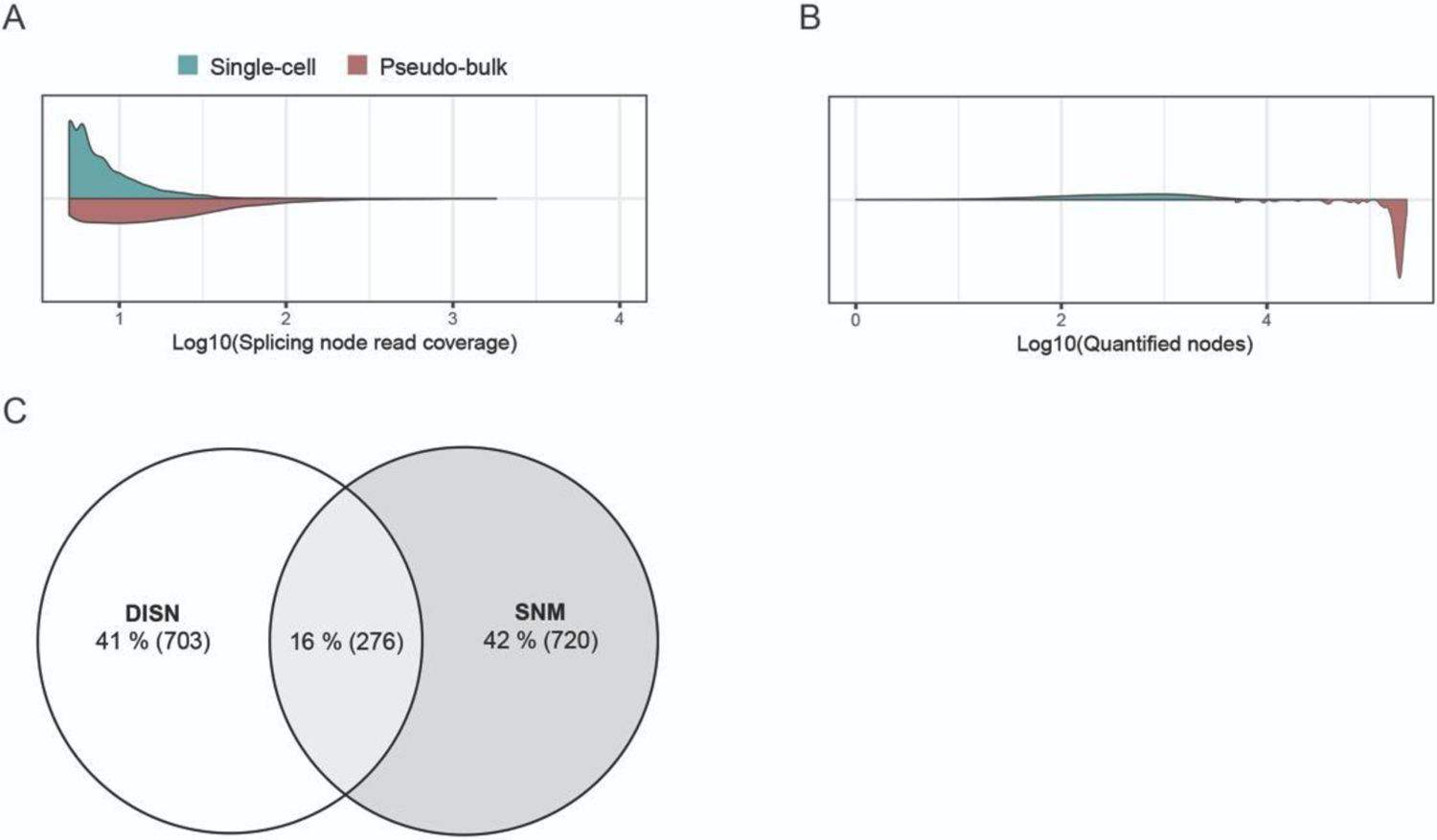
Global splicing marker analysis A. Violin plot showing the distribution of coverage values obtained for splice nodes when computed at single-cell or pseudo-bulk level. B. Violin plot showing the number quantified spliced nodes (read coverage>5) obtained when quantified at the single-cell or pseudo-bulk level. C. Euler diagram showing the splicing node intersection between the DISN and SNM sets.

**Supplementary Figure 5.**
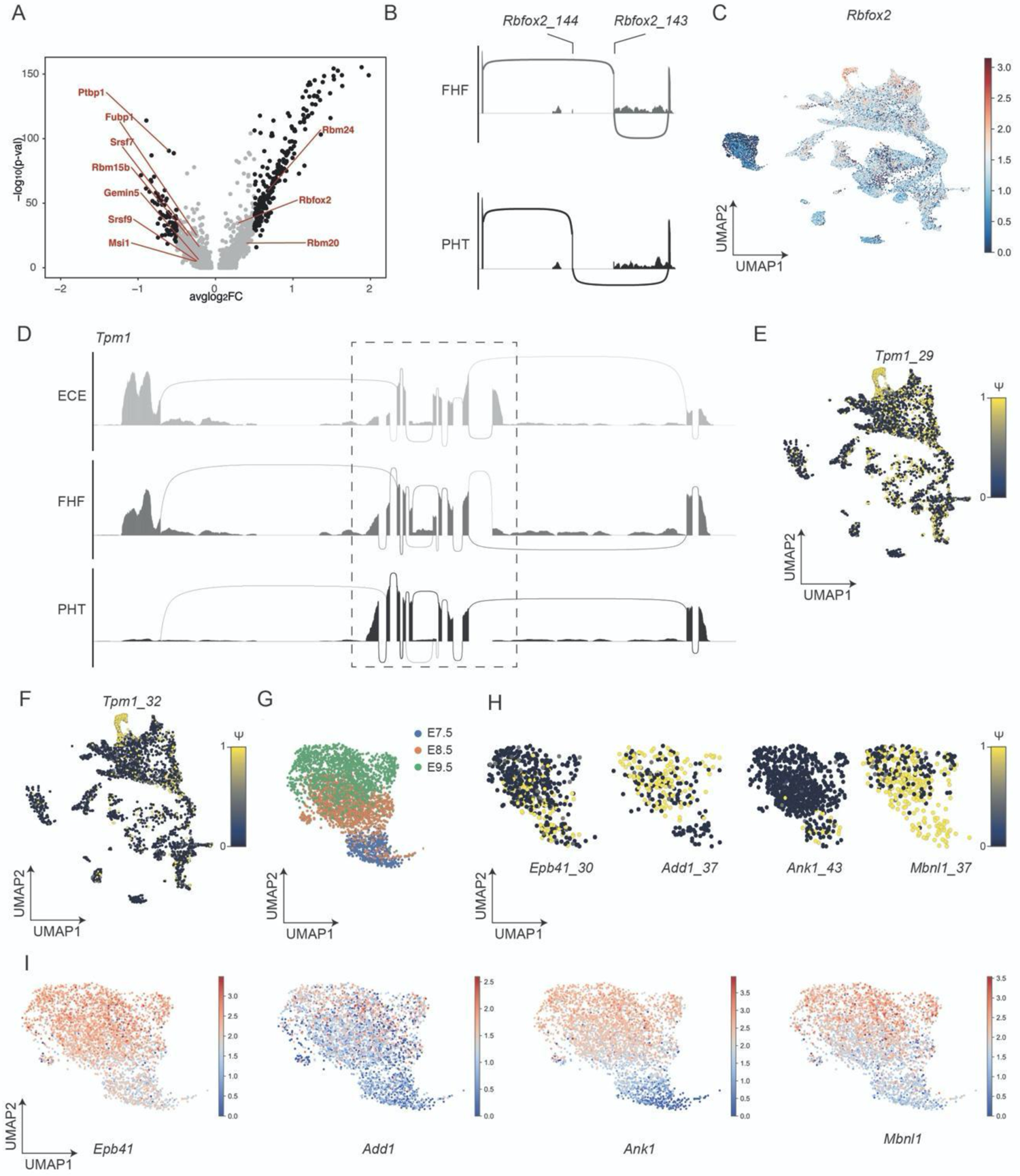
Heart and blood development reveal tissue-specific AS patterns across developmental trajectories A. Differential gene expression analysis using a Wilcoxon rank sum test between the FHF (negative average log2 fold-change values) and the PHT (positive average log2 fold-change values) with differentially expressed RBPs highlighted. Significance levels are indicated by color (grey non-significant and black significant), and determined by the following threshold: |average log2 fold-change | > 0.5 and Bonferroni adjusted p-value < 1E-05). B. *Rbfox2*_143 and 144 mutually exclusive exon usage in the FHF and PHT respectively. C. *Rbfox2* gene expression across the atlas, log2 normalized values. D. *Tpm1* sashimi plot between the ECE, FHF and PHT, dashed square highlights the region of interest plotted in Figure 7C. E. *Tpm1_29* single-cell PSI UMAP plot across the atlas highlighting a PHT specific core exon usage at the C-terminus. F. *Tpm1_32* single-cell PSI UMAP plot across the atlas highlighting a PHT specific core exon usage at the C-terminus. G. UMAP plot across timepoints depicting erythropoietic cell types. H. Single-cell PSI UMAPs of *Epb41_30*, *Add1_37*, *Ank1_43* and *Mbnl1_37* depicting alternative exonic usage across blood maturation trajectories. I. Single-cell gene expression UMAP plot depicting differences in gene expression for *Epb41*, *Add1*, *Ank1* and *Mbnl1* illustrating differences in gene expression that differ from the AS patterns observed across blood maturation.

**Supplementary tables are available at:** https://github.com/anna-alemany/vasaseq,

**Supplementary Table 1.** Lists of all equivalent cluster markers for protein coding (spliced and unspliced) genes and for lncRNA for each technology at different timepoints, detected using the default function from the scanpy package.

**Supplementary Table 2.** List of differentially expressed genes as detected using the t-test (absolute value of the log2 fold-change > 4; p-value below 0.001) between 10x and VASA-seq for each equivalent cluster in each timepoint, showing their mean expression, the standard deviation, and the fraction of cells expression the genes within the cluster. A pseudocount equal to the minimum mean expression value above zero was used to extract the log2 fold-change.

**Supplementary Table 3.** Cell-cycle genes during mouse embryonic development, obtained by differential gene expression analysis between S-phase and non-S-phase cells, for either pooled or separate timepoints.

**Supplementary Table 4.** Regressed data was clustered using the Leiden algorithm. Table showing differential expressed genes per cluster. These clusters were further used for cell type calling based on the differentially expressed genes (marker genes).

**Supplementary Table 5.** Differentially expressed histone genes between Leiden clusters/cell types. Both uniquely and multi-assigned histone genes were included.

**Supplementary Table 6.** Genes that contributed to the RNA velocity vector for VASA-seq and 10x. We found that the majority of significant genes were shared between the methods (1,492), but VASA-seq detected a large number of additional genes (1,069).

**Supplementary Table 7.** List of differentialy included nodes that were found across the different comparisons made between cell clusters. Leiden cluster IDs that were compared are indicated by A.cluster_names and B.cluster_names columns. Columns 4th to 8th provide the information corresponding to the assessed splicing nodes. Each comparison was repeated 50 times, and summary statistics (such as mean, standard deviation and variance) are reported for each splicing node that was differentially included across the computed comparisons between clusters. Finally, associated CDF beta values to each listed splicing node are indicated on the CDF beta column.

**Supplementary Table 8.** List of SNM identified across cell clusters. For each splicing node, the coordinate and the Leiden ID of the cluster, where they were found to be markers, are indicated by the Coord and Leiden columns, respectively. Additional stats that were used to identify each SNM are reported between columns 4th to 9th.

**Supplementary Table 9.** List of DISNs between the FHF and PHT, as identified by the MicroExonator pipeline. Positive DeltaPSI.mean values indicate inclusion in the FHF whereas negative values indicate inclusion in the PHT.

**Supplementary Table 10.** List of DISNs between the primitive erythroids at E7.5 and E9.5, as identified by the MicroExonator pipeline. Positive DeltaPSI.mean values indicate inclusion at E7.5 whereas negative values indicate inclusion at E9.5. Abs_psi represents the absolute value of the differential PSI.

**Supplementary Table 11.** List of oligonucleotides not found in cited publications. 195 rRNA depletion probes, two oligos for library prep of VASA-drop samples and 92 dual index PCR primers for amplification of VASA-drop samples.

## MATERIALS AND METHODS

### Ethics Statement

Experiments were performed in accordance with EU guidelines for the care and use of laboratory animals and under the authority of appropriate UK governmental legislation. Use of animals in this project was approved by the Animal Welfare and Ethical Review Body for the University of Cambridge and relevant Home Office licence PPL (7677788) is in place.

### Cell Lines

HEK293Ts were passaged every second day and cultured in T75 flasks. The culture media was DMEM-F12 (ThermoFisher) supplemented with 10% heat-inactivated FBS (ThermoFisher) and 100 U/ml Penicillin-Streptomycin (ThermoFisher). For passaging, the cells were washed with 10 ml of ice-cold 1x PBS (Lonza) twice. 9 ml of PBS was added to the flask and cells were detached by adding 1 ml of 10xTrypsin-EDTA (Sigma-Aldrich) and incubated at 37 °C for 5 minutes. Trypsin-EDTA was then inactivated with 15 ml of DMEM-F12 + 10% FBS and incubated at 37 °C for 5 minutes. The cells were then pelleted at 300g for 3 minutes and the supernatant was aspirated. After aspiration of the supernatant, the cells were washed twice in PBS and viability-assessed and counted before encapsulation.

Mouse embryonic stem cells (mESCs) were passaged every other day and cultured in 2i+LIF medium. Briefly, Dulbecco’s Modified Eagle Medium F-12 (DMEM/F-12) Nutrient Mixture, without L-Glutamine (ThermoFisher Scientific, 21331020) and Neurobasal Medium without L-Glutamine (ThermoFisher Scientific) in a 1:1 ratio, 0.1 % Sodium Bicarbonate (ThermoFisher Scientific), 0.11 % Bovine Albumin Fraction V Solution (ThermoFisher Scientific), 0.5x B-27 Supplement (ThermoFisher Scientific, 17504044), 1x N-2 Supplement (Cambridge Stem Cell Institute made in house), 50 µM 2-Mercaptoethanol (ThermoFisher Scientific), 2 mM L-Glutamine (ThermoFisher Scientific), 100 U/ml Penicillin-Streptomycin (ThermoFisher Scientific), 12.5 µg/ml Insulin Zinc (ThermoFisher Scientific), 0.2 µg/ml mLIF (Cambridge Stem Cell Institute), 3 µM CHIR99021 (Cambridge Stem Cell Institute), 1 µM PD0325901 ( Cambridge Stem Cell Institute). Culture dishes were coated with 0.1 % gelatine in phosphate-buffered saline (PBS) for at least 30 min. Cells were detached with 500 μl/6-well Accutase (Merck) for 3 minutes at 37 °C. The detached cells were transferred into 9.5 ml washing medium (DMEM/F-12 with 1% Bovine Albumin Fraction V Solution) and centrifuged at 300 g for 3 minutes. The supernatant was aspirated and the cell pellet was resuspended in 2i/LIF medium and re-plated at 80,000 cells/6-well. For the encapsulation process, the cells were washed twice in PBS, viability-assessed and counted before dilution to the correct concentration.

### Murine embryo collection and dissociation

Pregnant C57BL/6 female mice were purchased from Charles River or obtained from natural mating of C57BL/6 mice in house. Mice were maintained on a lighting regime of 12:12 hours light:dark with food and water supplied ad libitum. Detection of a copulation plug following natural mating indicated embryonic day (E) 0.5. Following euthanasia of the females using cervical dislocation, the *uteri* were collected into PBS (Lonza) with 2% heat-inactivated FBS (Gibco, Thermo Fisher Scientific) and the embryos were immediately dissected and processed for scRNA-seq. Mouse embryos were dissected at time points E6.5, E7.5, E8.5 and E9.5 as previously reported (PMID: 8269852). Embryos from the same stage were pooled ina LoBind tube (Eppendorf). E8.5 and E9.5 embryos were cut into pieces under a stereomicroscopy before collecting into a tube. The pooled samples were centrifuged at 300g for 5 min. The supernatant was aspirated and 100-200 µl of TrypLE Express (Gibco) dissociation reagent was added to the samples. The tube was incubated at 37 °C for a minimum of 7 minutes (or until completely dissociated) in an orbital shaker. Subsequently, 1 ml FBS was added to the tube to inactivate TrypLE. The sample was repeatedly centrifuged and washed with PBS before finally being resuspended in PBS supplemented with 0.04% BSA, and filtered through a 40 µm Flowmi Tip Strainer (ThermoFisher Scientific).

### VASA-plate: Cell sorting in 384-plates

Single cells were sorted into 384-well hardshell plates (BioRad) using a BD FACSJazz. Each well was pre-filled with 5 µl mineral oil (Sigma-Aldrich) and 50 nl Cel-seq2/Sort-seq (Hashimshony et al., 2012; Muraro et al., 2016) primer with a concentration of 0.25 µM. Plates were sealed (Greiner, Silverseal sealer, 676090) and spun down at 2,000 rcf for 1 minute (Eppendorf 5810R) before stored at −80 °C.

### VASA-plate: Cell lysis and RNA fragmentation

All dispensions were carried out with a Nanodrop II (Innovadyne Technoligies Inc.), all incubations with a GeneAmp PCR System 9700 Thermocycler (Applied Biosystems) and all spinning steps with an Eppendorf 5810R unless otherwise specified.

50 nl of lysis and fragmentation mix (3.4x First Strand Buffer (Invitrogen), 1.2 mUnits Thermolabile Proteinase K (NEB) and 0.2 % IGEPAL CA-630 (Sigma-Aldrich)) was added to each well. Plates were sealed and spun down at 2,000 rcf for 2 min. Lysis was carried out at 25 °C for 1 hour followed by 55 °C at 10 minutes. Plates were snapp chilled on ice, before fragmentation was carried out at 85 °C for 3 minutes. Plates were snapp chilled, spun down at 2,000 rcf for 1 minute and stored on ice before next dispensation.

### VASA-plate: RNA repair and poly(A)-tailing

50 nl of RNA repair and poly(A)-tailing mix (0.6x First Strand Buffer (Invitrogen), 20 mM DTT (Invitrogen), 7.5 nM ATP (NEB), 37.5 mU *E.coli* Poly(A) Polymerase (NEB), 50 mU T4 PNK (NEB) and 10 mM RNaseOUT (Invitrogen)) was added to each well. Plates were sealed and spun down at 2,000 rcf for 2 minutes. Repair and tailing was carried out at 37 °C for 1 hour. Plates were snapp chilled, spun down at 2,000 rcf for 1 minute and stored on ice before next dispensation.

### VASA-plate: Reverse transcription

50 nl of reverse transcription mix (2 mM (each) dNTP mix (Promega) and 0.8 Units Superscript III (Invitrogen)) was added to each well. Plates were sealed and spun down at 2,000 rcf for 2 minutes. Reverse transcription was carried out at 50 °C for 1 hour. Plates were snapp chilled, spun down at 2,000 rcf for 1 minute and stored on ice before next dispensation.

### VASA-plate: Second strand synthesis

1100 nl of Second strand synthesis mix (1.14x Second Strand Buffer (Invitrogen), 0.23 mM (each) dNTP mix (Promega), 0.35 Units *E. coli* DNA Polymerase I (Invitrogen) and 20 mU RNaseH (Invitrogen)) was added to each well. Plates were sealed and spun down at 2,000 rcf for 2 minutes. Second strand synthesis was carried out at 16 °C for 2 hours followed by 85 °C for 20 minutes. Plates were snapp chilled, spun down at 2,000 rcf for 1 minute and stored on ice before pooling. The protocol for pooling and IVT was the same as Sort-seq (Muraro et al., 2016).

### VASA-drop: Design of the droplet generation device

The droplet generation device for compressible barcoded bead and single-cell co-encapsulation (Figure S1B) was modified from previous designs (Klein et al., 2015; Zillionis et al., 2017). The flow-focusing junction (80 µm) was narrowed to generate smaller droplets (0.55 nl) at high-throughput (300 Hz).

### VASA-drop: Design of droplet pico-injection devices

The design of both droplet pico-injector devices is based on the findings of a previous study (Abate et al., 2010). Several key features were added to the architecture of previous designs to ameliorate the robustness of the injections in large droplets containing compressible barcoded beads:

1. A diluting oil channel, number 3 (Figure S1B), that reduces the packing of the emulsion to eliminate fragmentation of densely packed droplets before being re-injected in the pico-injection channel. This design feature allows for packed droplets to arrange into an evenly-spaced monolayer that reduces fluctuations in volume of droplets after pico-injection.
2. Smooth narrowing of the reinjection chamber facilitating the ordering of droplets before spacing which reduced droplet break-up.
3. Deepening of the outlet junction before the outlet (Figures S1C and S1D, deep blue color), which stabilizes droplets and reduces droplet merging, which was observed during the rapid transition from the shallow microfluidic channel to a wide tubing or collection tip.

### VASA-drop: Photolithography of microfluidic moulds

The microfluidic devices were fabricated following standard hard lithography protocols. First, microfluidic moulds were patterned on 3” size silicon wafer (Microchemicals) using high-resolution film masks (Microlithography Services Ltd) and SU-8 2075 photoresists (Kayaku Advanced Materials). A MJB4 mask aligner (SUSS MicroTec) was used to UV expose all the SU8 spin-coated wafers. The thickness of the channels was measured using a DektakXT Stylus profilometer (Bruker).

We used the following settings for photolithography:

|  | Fabrication step (no. of layer) |  |
| --- | --- | --- |
|  | 1<sup>st</sup> layer | 2<sup>nd</sup> layer (used only for pico-injectors) |
| Nominal thickness | 80 $\mu\text{m}$ | 80$\mu\text{m}$ 2<sup>nd</sup> layer (160 $\mu\text{m}$ final thickness) |
| Resist used | SU8-2075 | SU8-2075 |
| Spin coating speed | 1st step: 10 sec, 500rpm<br>2nd step: 30 sec, 2750 rpm | 1st step: 10 sec, 500rpm<br>2nd step: 30 sec, 2750 rpm |
| Pre-baking | 3 min at 65°C<br>9 min at 95°C | 3 min at 65°C<br>9 min at 95°C |
| Exposure (at $\sim 10 \text{ mW cm}^2$) | 2 x 10 sec | 2 x 10 sec |
| Post-baking | 2 min at 65°C<br>7 min at 95°C | 2 min at 65°C<br>7 min at 95°C |
| Development in the beaker filled with 30-50ml of PGMEA (Propylene glycol methyl ether acetate, Sigma-Aldrich) | Approximately 5 minutes until all uncured SU8 is removed from the wafer, development time depends on the intensity of manual agitation (only for a 1-layer chip) | Approximately 10 minutes until all uncured SU8 is removed from the wafer, development time depends on the intensity of manual agitation |
| Hard baking (optional) | 10 min at 200 °C (only for a 1-layer chip) | 10 min at 200 °C |
| Measured range of thicknesses | 80-84 $\mu\text{m}$ | 168-178 $\mu\text{m}$ (second layer is usually ca. 20% thicker than nominal) |

### VASA-drop: Soft lithography

To manufacture PDMS microfluidic devices, 20-30 grams of silicone elastomer base and curing agent (Sylgard 184, Dow Corning) were mixed at a 10:1 (w/w) ratio in a plastic cup and degassed in a vacuum chamber for 30 minutes. PDMS was then poured on a SU8 master wafer and cured in the oven at 65 °C for at least 4 hours. Next, the inlet holes were punched using 2 types of biopsy punchers with plungers (Kai Industries): a 1.5mm diameter punch was used to make the inlet for cell delivery tip, number 2 (Figure S1B), outlet tip for droplet collection, number 4 (Figure S1B) and number 5 (Figure S1C) and the inlet for droplet reinjection, number 1 (Figures S1C and S1D), while other inlets were made using a 1mm wide biopsy puncher. The patterned PDMS chip was then plasma bonded to a 52 mm x 76 mm x 1 mm (length x width x thickness) glass slide (VWR) in a low-pressure oxygen plasma generator (Femto, Diener Electronics). Next, the hydrophobic modification of microfluidic channels was performed by flushing the device with 1% (v/v) trichloro(1H,1H,2H,2H-perfluorooctyl)-silane (Sigma-Aldrich) in HFE-7500 (3M) and baked on a hot plate at 75 °C for at least 30 minutes to evaporate the fluorocarbon oil and silane mix.

### VASA-drop: Cell loading and droplet collection/re-injection chamber manufacturing

A) Cell injection chamber

The cells were loaded in a cell loading tip pre-filled with mineral oil (Sigma-Aldrich). To manufacture the tip, a low-retention tip with 200 µl volume capacity (Axygen) was cut at the top, in parallel to the rim and under the filter. A solidified 3-mm thick piece of PDMS (Dow Corning) was punched with a 5.0 mm sampling tool (EMS-core). The circular piece of PDMS was then biopsy punched with a 1mm wide biopsy puncher (Kai Industries) in the middle. The circular piece of PDMS was pushed inside the tip, while remaining parallel to the cut. A 1 ml glass syringe (SGE) was then pre-filled with 1 ml of mineral oil and connected to a 30 cm long tubing (Portex) that can be connected to the inserted PDMS. The tip was then pre-filled with mineral oil by manually pushing the syringe, and the cell-containing solution was further aspirated with care, as to not introduce any air bubble in the system. The tip can then be connected to the cell-encapsulation PDMS device, number 2 (Figure S1B), and injection rates are modulated by a Nemesys syringe pusher (Cetoni).

B) Droplet collection and reinjection chamber

A second type of tip chamber was designed to collect, incubate and reinject droplets for each microfluidic step. To this end, A 5mm-thick PDMS piece was punched with a 8.0 sampling tool (EMS-core), and re-punched in its center using a 1mm wide biopsy puncher (Kai Industries) and a 30cm long tubing (Portex) was connected to the latter punched hole. The resulting piece of PDMS was then inserted into a 1 ml filterless pipette tip (Sigma-Aldrich), with a parallel orientation to the rim. Unsolidified PDMS (Dow Corning, 1:10 (w/w) ratio, degassed) was then inserted into the space between the rim and the circular PDMS piece at the top. The tips were then incubated at 65°C for at least four hours and connected to a 1 ml glass syringe (SGE) pre-filled with mineral oil that can be pushed using a neMesys syringe pusher (Cetoni) to re-inject droplets into the pico-injectors, number 3 (Figures S1C and S1D). To collect the droplets after the initial encapsulation or at the end of the first pico-injection, the tip can be connected to the outlet of the devices, number 5 (Figures S1C and S1D), and the syringe is disconnected to allow the evacuation of mineral oil as the tip gets loaded. For each of the re-injection and collection steps, the PDMS-punched holes on the microfluidic device need to be primed with 5% (w/w) RAN 008-fluorosurfactant (RAN biotechnologies) in HFE7500 (3M) as to avoid a trapped air bubble to perturbate the stability of re-injection or the integrity of the emulsions (Figure S1E). The tip can be closed by the narrower end by punching a 8 mm-thick piece of PDMS with a 1.5 mm biopsy buncher that was bonded to microscopy glass using an oxygen plasma, which closes the system and allows for incubation of the tip in the water bath.

### VASA-drop: Microfluidic device operation

A) Polyacrylamide beads manufacturing

Barcoded polyacrylamide beads were manufactured following the protocol previously described (Zilionis et al., 2017). Briefly, a polyacrylamide mix was compartmentalized in 60 µm water-in-oil emulsions using a single-inlet flow-focusing device and collected in a 1.5 ml LoBind tube (Eppendorf) containing 200 µl of mineral oil (Sigma-Aldrich). The droplets were solidified overnight at 65°C, de-emulsified using 20% 1H,1H,2H,2H-Perfluoro-1-octanol (Alfa Aesar) in HFE7500 (3M) and stored at 4°C for up to 6 months.

B) Co-encapsulation of cells and barcoded beads

A detailed protocol (Zilionis et al., 2017) was used as a reference for droplet generation. First, the microfluidic droplet generation chip was installed on the stage of an inverted microscope (Olympus XI73). Next, two pieces of polyethylene tubing (Portex) were connected to two 1 mL gas-tight syringes (SGE) and filled with sterile PBS (Lonza) buffer. The tubing was manually filled with PBS and a small, 1-cm long, bubble was left at the end tip of each tubing. The bead suspension and lysis mix were manually aspirated to the tubing and the small bubble provided a separation between the reagents and the PBS buffer. 150 µl of cell suspension was manually aspirated into the cell chamber pre-filled with mineral oil (Sigma-Aldrich). A fourth 2.5 ml glass syringe (SGE) was filled with 5% (w/w) 008-fluorosurfactant (RAN Biotechnologies) in HFE-7500 (3M). Next, all three tubings and the cell chamber with cell suspension were inserted to the corresponding inlets of the droplet generation chip (Fig. S1A).

Four neMesys syringe pumps (Cetoni) were used to flow each component and the droplet formation was monitored using 4x or 10x objectives and a fast camera (Phantom) connected to the inverted microscope. After the device was primed and droplet generation was stabilized, the collection chamber was connected to the outlet.

C) Microfluidic device operation – picoinjection

Before starting the pico-injection of droplets containing single-cell lysates, the electrode section, number 6&7 (Figures S1C and S1D), of the device were prefilled with filtered 5M NaCl as previously described (Sciambi et al., 2014). 5% (w/w) 008-fluorosurfactant (RAN Biotechnologies) in HFE-7500 (3M) using a pre-filled 2.5 ml glass syringe (SGE) connected to a piece of tubing (Portex). The reaction mix was primed and the tip containing the emulsion (with fluorinated oil evacuated by pushing the glass syringe until the emulsions reached the exit of the tip) were primed and connected to the device. Next, the flows of emulsion, mix and two streams of oil, diluting: number 3 (Figure S1C), spacing: number 3 (Figure S1C), were applied using the neMesys syringe pumps. The droplets were diluted in a first instance in the re-injection chamber and then spaced with the second stream of oil in a flow-focusing reinjection junction with 5% (w/w) 008-fluorosurfactant (RAN Biotechnologies) in HFE-7500 (3M). The function generator (Thurlby Thandar Instruments) was set to generate square waves of 2.5 V amplitude and 10 kHz frequency which was further amplified 100 times to 250V by a Trek 601C-1 amplifier which enabled coalescence-activated droplet injection of the reagent into the droplets. The droplets were collected in a 1 ml collection tip connected at the outlet, number 5 (Figure S1C).

### VASA-drop: Polyacrylamide bead barcoding

The bead barcoding procedure was performed as previously described (Zilionis et al., 2017), but using the inDrop v3 barcoding scheme (Briggs et al., 2018). Briefly, the solidified barcoded beads were filtered and dispensed in four 96-well plates containing the first barcode from the inDrop v3 design, which was annealed and the strand extending from the bead-bound adapter was added using a Bst 2.0 DNA polymerase (NEB). The reaction was then stopped, and the second strand was removed using a sodium hydroxide treatment. The second barcode was added in a similar fashion, and the beads were stored up to six months at 4°C.

### VASA-drop: Cell encapsulation in water-in-oil emulsions

For the cultured cells and the embryos, we used a loading concentration of 450 cells/µl in 1xPBS (Lonza) with 15% Optiprep (Sigma-Aldrich). The lysis mix was made fresh before each encapsulation, as follows: 0.5 mM dNTPs each (ThermoFisher), 0.52% IGEPAL-CA630 (Sigma-Aldrich), 40 mM Ultrapure Tris-HCl pH 8 (Life sciences), 3.76x First Strand Buffer (Invitrogen), 3 mM Magnesium Chloride (Ambion) and 6 U/ml Thermolabile proteinase K (NEB). The barcoded PAAm beads were prepared for encapsulation as previously described (Zilionis et al., 2017). The lysis mix and bead suspensions were loaded in the tubing of two individual 1 ml SGE glass syringes filled with PBS (Lonza), and separated by an air bubble from the reagents in the tubing as previously described. The cells were loaded in a cell injection container pre-filled with mineral oil (Sigma-Aldrich) as previously described. The injection flow rates for the droplet encapsulation device (Figure S1B) were: 1) the cell suspension was flown at 85 µl/hr, number 2 (Figure S1B), the bead suspension was flown at 65 µl/hr, number 3 (Figure S1B), the lysis solution was flown at 75 µl/hr, number 1 (Figure S1D), and 5% (w/w) 008-fluorosurfactant (RAN Biotechnologies) in HFE-7500 (3M) surfactant was flown at 450 µl/hr using a 2.5 ml glass syringe (SGE), number 4 (Figure S1B). All flow rates for each microfluidic manipulation were generated using neMesys pumps (cetoni). The average droplet size was ∼0.55 nl for these flow rates and a microfluidic device depth of 60 µm. The droplets were collected for approximately one hour in a 1 ml pipette tip (Greiner) pre-filled with mineral oil at the outlet, number 5 (Figure S1B) and connected to a tubing via a PDMS connector. The collection tip was then closed using a 1ml SGE glass syringe pre-filled with mineral oil and connected to a glass-bonded PDMS plug.

### VASA-drop: Cell lysis and RNA fragmentation

The tip container was further left at room temperature (23°C) for 20 minutes to allow for cell lysis to occur and the tip was further placed under a High-Intensity UV Inspection Lamp (UVP) that was switched on for 7 minutes for barcode photocleavage. The container was then further submerged in a water bath (Grant JB) placed at 85°C for 6 minutes and 30 seconds. The container was then immediately submerged in an ice bucket filled up with half proportions of ice and water to stop the reaction.

### VASA-drop: First pico-injection for RNA repair and poly(A)-tailing

The droplets were re-injected in the first pico-injector device with the shorter re-injection channel (Figure S1C) and coalescence-induced merging with a poly(A) solution consisting of 26.6 mM Tris-HCl pH 8 (Invitrogen), 15.8 mM DTT (Invitrogen), 0.83x First Strand buffer (Invitrogen), 0.19 mM ATP (NEB), 3.15 kU/ml T4 Polynucleotide kinase (NEB), 250 U/ml *E. coli* poly(A) polymerase, 2.6 kU/ml RNaseOUT (Applied biosystems). The merging was applied by prefilling the electrode section, number 6&7 (Figure S1B) of the device with 5M NaCl as previously described (Sciambi et al. 2014). The function generator was set to generate square waves of Amplitude 2.5 V and 10 kHz frequency. The droplets were spaced in a first instance before re-injection in a flow-focusing junction using 5% (w/w) 008-fluorosurfactant (RAN Biotechnologies) in HFE-7500 (3M). The flow rates used were 200 µl/hr for the droplets, number 1 (Figure S1C), 60 µl/hr for the poly(A) mix, number 4 (Figure S1C), 120 µl/hr for the first spacing oil, number 2 (Figure S1C) and 400 µl/hr for the second spacing oil, number 3 (Figure S1C). This generated ∼0.8 nl droplets at 70Hz. The droplets were collected in a 1 ml collection tip (Greiner) inserted to the outlet, number 5 (Figure S1C), and incubated for 25 minutes at room temperature (23°C) followed by 8 minutes in a 37°C water bath (Grant JB). The collection tip was then closed at each end by a PDMS plus and connecting a 1ml glass syringe (SGE) pre-filled with mineral oil (Sigma-Aldrich) and submerged in an ice-cold water bath for 2 minutes. The droplets were immediately processed for reverse-transcription after that.

### VASA-drop: Second pico-injection for reverse transcription

The droplets were re-injected in the second pico-injector (Figure S1D) similarly to the previous step, albeit the injected droplets were collected in fractions of ∼1000 cells (∼27 µl of loaded droplets) in 1 ml LoBind tubes (Eppendorf) pre-filled with 200 µl of mineral oil. The droplets were injected with a reverse transcription mix constituted of 25 mM Tris-HCl pH 8 (Invitrogen), 8 mM DTT (Invitrogen), 0.75x First Strand buffer (Invitrogen), 1 mM dNTPs, 20 kU/ml Superscript III (Invitrogen), 1.2 kU/ml RNAseOUT (Applied biosystems). The flow rates for this device were as follows: 70 µl/hr for the first spacing oil, number 2 (Figure S1D), 700 µl/hr for the second spacing oil, number 3 (Figure S1D), 300 µl/hr for the re-injected droplets, number 1 (Figure S1D) and 255 µl/hr for the RT mix, number 4 (Figure S1D). The collected fractions were incubated at 50°C for 2 hours and then heat-inactivated at 70°C for 20 minutes. For de-emulsification of the droplets, the mineral oil and oil phase were aspirated and discarded. Then 200 µl of filtered HFE7500 was added to the emulsions, followed by 200 µl of 100% 1H,1H,2H,2H-Perfluoro-1-octanol. The tubes were centrifuged for 5 seconds on a tabletop centrifuge, then 300 µl of the oil phase was removed and 100 µl of fresh HFE7500 oil was added, as well as 50 µl of TE buffer (Zymo). At this point the fractions were stored at − 80°C. The protocol up to and including the IVT step was the same as for inDrop (Zilionis et al., 2017).

### VASA-plate & VASA-drop: Downstream library preparation and sequencing

For VASA-plate: after IVT, 2 µl ExoSAP-IT (Applied Biosystems) was added, and each sample was incubated at 37°C for 15 minutes. A 1.8x volumetric ratio AmpureXP clean-up was then performed and the aRNA was eluted in 10 µl of nuclease-free water. The purified aRNA’s concentration was measured using a Qubit (Invitrogen), and the concentration adjusted to max 100 ng/µl. 6 µl per sample was mixed with 2 µl of rRNA depletion probes (25 µM) (reverse complement of published probes (Adiconis et al., 2013)) and 2 µl of Hybridization buffer (pH 7.5, 500mM Tris-HCl, 1M NaCl). Samples were incubated at 95°C for 2 minutes and brought to 45°C with a gradient of 0.1°C/s. Once the probes were hybridised, 2 µl of Thermostable RNAseH (Epicentre) and 8 µl of RNAseH buffer (pH 7.5, 125mM Tris-HCl, 250mM NaCl, 50mM MgCl2) was added. The reaction was incubated at 45°C for 30 minutes and further kept on ice. Next, 4 µl of RQ DNAse I (Promega), 21 µl of nuclease-free water and 5 µl of CaCl_2_ (10mM) was added to the reaction mixture. The mixture was further incubated at 37°C for 30 minutes, followed by snap cooling on ice. A 1.6x volumetric ratio AmpureXP clean-up was then performed and the aRNA was eluted in 6 µl of nuclease-free water. Next, 1 µl of RA3 ligation oligonucleotide (20 µM, Table S11) was added to 5 µl of the aRNA, and the reaction was brought to 70°C for 2 minutes, followed by snap cooling on ice. This was followed by the addition of 1 µl 10x T4 RNA ligase reaction buffer (NEB), 1 µl NEB T4 RNA Ligase2, truncated (NEB), 1 µl RNAseOUT (Invitrogen) and 1 µl nuclease-free water, The reaction was incubated at 25°C for 1 hour, followed by snap cooling on ice. The adapter ligated aRNA was then mixed with 1 µl of dNTPs (10 mM each) (Promega) and 2 µl of RTP oligonucleotide (20 µM, Table S11). The mixture was incubated at 65°C for 5 minutes, followed by snap cooling on ice. Next, 4 µl 5x First strand synthesis buffer (Invitrogen), 1 µl nuclease-free water, 1 µl 0.1M DTT (Invitrogen), 1 µl RNAseOUT and 1 µl Superscript III was added to the sample. The reaction was incubated at 50°C for 1 hour, followed by 70°C for 15 minutes and then snap cooled on ice. To reduce excess RNA material, 1 µl of RNAseA (ThermoFisher) was further added to each tube and the cDNA was incubated at 37°C for 30 minutes, followed by a 1x volumetric AmpureXP clean-up. The cDNA was eluted in 20 µl of nuclease-free water. Half the material was used for the final PCR (10 µl). Each sample was mixed with 25 µl NEBNext High-Fidelity 2X PCR Master Mix (VASA-plate) or Kapa HiFi HotStart PCR mix (VASA-drop), 4 µl PE1/PE2 primer mix (5 μM each) (Hashimshony et al., 2012; Muraro et al., 2016) (VASA-plate) or 5 µl PE1/PE2 primer mix (5 μM each) (Table S11) (VASA-drop) and 11 µl (VASA-plate) or 10 µl (VASA-drop) of nuclease-free water. The samples were amplified with the following PCR programs: VASA-plate: Initial heat denaturation for 30 seconds at 98°C, 7-8 cycles for 10 seconds at 98°C, 30 seconds at 60°C, 30 seconds at 72°C, and final extension for 10 minutes at 72°C. VASA-drop: Initial heat denaturation for 2 minutes at 98°C, 2 cycles for 20 seconds at 98°C, 30 seconds at 55°C, 40 seconds at 72°C, 5-6 cycles for 20 seconds at 98°C, 30 seconds at 65°C, 40 seconds at 72°C and final extension for 5 minutes at 72°C. Each amplified and indexed sample was purified twice using a 0.8x volumetric ratio of AmpureXP beads and eluted in 10 µl. Final libraries were checked for proper length on a Bioanalyzer (Agilent) and concentration was measured with a Qubit (Invitrogen).

The VASA-drop samples were sequenced on a Novaseq 6000 S2, 300 cycles flow cell (Illumina), with the following parameters: Read1 247 cycles, Index1 31 cycles, Index2 8 cycles, Read2 14 cycles. VASA-plate samples were sequenced on a NextSeq 500, High output 150 cycles flow cell (Illumina), with the following parameters: Read1 26 cycles, Index 8 cycles, Read2 135 cycles.

### Fastq file preprocessing in VASA-drop and 10X

Raw reads for VASA-drop were pre-processed with a python script to have a favorable format for the pipeline (four reads were demultiplexed and rearranged into two reads). For each biological read (read 1), the UMI (6-nt long in VASA, 10-nt long in 10x) and the cell-specific barcode (16-nt long in VASA, 14-nt long in 10x) were extracted. To determine the number of cells in each sample, first the total number of raw reads was determined for each possible barcode. Next, we plotted the histogram of log10 (read number) for each possible barcode, which we fitted to a polynomial function that shows 2 or 3 minima. We used the position of the minimum with the highest value of log10(reads) as the threshold: only barcodes with reads above this threshold are used for downstream analysis. We merged sequenced barcodes that can be uniquely assigned to an accepted barcode with a Hamming distance of 2 nucleotides or less.

### Fastq file preprocessing in VASA-plate

Read 1 starts with 6-nt long UFI/UMI, followed by an 8-nt long cell-specific barcode. There are only 384 cell-specific barcodes, each one corresponding to a well in a 384-well plate (available in GSE176588). We merged sequenced barcodes that can be uniquely assigned to an accepted barcode with a Hamming distance of 1 nucleotide or less.

### Mapping data (VASAseq, 10x, Smart-seq v3)

Read 2 was assigned to accepted barcodes (extracted from read 1) were trimmed with TrimGalore (v. 0.4.3) with default parameters. Next, homopolymers at the end of the read were removed with cutadapt (v. 2.10).

*In silico* ribosomal depletion was performed by mapping the trimmed reads to mouse or human ribosomal RNA (ncbi) using bwa mem and bwa aln (v. 0.7.10). Multi- and single-mappers were filtered out. The remaining reads were mapped to the mouse mm10 genome or to the human GRCh38 genome (ensembl 99) using STAR with default parameters. Assignment of reads to gene biotypes was performed according to the following hierarchy:

– All mappings falling in TEC transcripts were discarded.
– Reads fully falling inside a region annotated as miscRNA, mtRNA, mttRNA, TrJGene, miRNA, rRNA, ribozymes, sRNA, scaRNA, snRNA or snoRNA (e.g, biotypes that do not have annotated introns) were assigned to such regions.
– When a read maps to multiple genes simultaneously (because of annotation overlap in the reference gtf file), exonic annotations were given preference to introns. In case all references are exonic or intronic, the read is assigned to a gene whose name is the sequence of all the target gene names.
– Reads falling into exon/intron junctions or inside introns are assigned to unspliced transcripts. Reads falling inside exonic regions are assigned to spliced transcripts.
– If at least one UFI of the same cell from the same transcript has been assigned to an unspliced transcripts (because if is mapped in an intron or an intron/exon junction), all the other reads with the same UFI of the same cell for the same transcript are automatically assigned to unspliced transcripts even if they mapped to exons exclusively.

### Benchmarking against other methods

To determine the number of potential doublets, barcodes with more that 75% of the genes assigned to only one of either mouse or human were considered singlets. Cells with less than 7,500 UFIs were filtered out and not assigned to any organism. For gene body coverage, the bam-files for all single cells were used as a bulk. QoRTs (Hartley and Mullikin, 2015) were used to calculate coverage, and only protein coding genes were kept. For Smart-seq3, both reads containing a UMI (5’ reads) and non-UMI containing reads were used together. Average coverages were used for the plotting. To determine percentages of different biotypes, all single cells were used as a bulk. UMI/UFI filtering was carried out for reads where this was possible. For Smart-seq3, both reads containing a UMI (5’ reads) and non-UMI containing reads were used together. For the gene detection assay, only cells that had been sequenced to the highest numbers of reads (reads with proper barcode and quality/homopolymers trimming) were used (75,000 for saturation curve and 750,000 deep sequencing comparison). For Smart-seq3, four cells, with much lower reads than the rest, were removed as they were considered failed libraries. Downsampling was carried out with DropletUtils (Griffiths et al., 2018; Lun et al., 2019) on the count matrices (non UMI/UFI filtered), based on the number of input reads and target reads, and only uniquely assigned genes were counted. For the percentage of intronic reads, each cell was used individually. UMI/UFI filtering was carried out for reads where this was possible. For Smart-seq3, both reads containing a UMI (5’ reads) and non-UMI containing reads were used together. Mean and standard deviation was calculated and plotted.

### scRNA seq analysis for mouse VASA-seq libraries and individual time points

The scrublet and scanpy packages were used together with custom-made code. In brief, for VASA seq only cells with more than 10^4^ (E6.5), 10^3.5^ (E7.5, E8.5), and 10^3^ (E9.5) reads and less than 10^6^ transcripts were kept. Next, only cells in which 85%-95% of transcripts belonging to protein coding genes, 1%-3% of transcripts belonging to lncRNA and 5%-15% of transcripts belonging to small RNA were kept. Unspliced and spliced protein coding genes were treated as different entries in our count tables to recover extra granularity in the downstream two-dimensional projection. Potential doublets as detected by scrublet with default parameters were removed. The resulting count tables were library-size normalized to 10^4^ transcripts and data was log-transformed with a pseudocount equal to 1. Cells with a total transcript count to histone genes above 35 were assumed to be in S-phase (Figure 4). Differential gene expression analysis between cells in S-phase and not S-phase was performed using the t-test to determine cell cycle genes (default scanpy.tl.rank_genes_groups function in scanpy), for separate timepoints and all data together (Table S3). Next, highly variable genes with mean log expression between 0.0125 and 5 are selected, and cell cycle genes were excluded. Number of counts and cell-cycle properties were regressed out (scanpy function scanpy.pp.regress.out), data was z-transformed (scanpy.pp.scale). For all timepoints, we selected the top 50 principal components (except for E6.5, for which we selected the first 20). For each timepoint, we constructed a directed graph connecting nearest neighbor cells in the reduced PCA space, using the Manhattan metric as described previously (Wanger et al., 2018). Initially, for each cell we identified its 10 nearest neighbors. An outgoing edge from cell *i* to cell *j* was kept if the distance *d_ij_* was less than the mean+1.5*standard deviation among all the distances connecting 10 nearest neighbors. Cells that are not connected to any other cell were filtered out. The directed graph was converted to an undirected graph and a two-dimensional Uniform Manifold Approximation and Projection (UMAP) was obtained as previously described (McInnes et al., 2020). We clustered the data using the Leiden algorithm (scanpy.tl.leiden, resolution set to 1), and performed differential gene expression between Leiden clusters using the t-test (default scanpy.tl.rangk_genes_groups).

### scRNA seq analysis for mouse 10x-seq libraries and individual time points

10x data was analyzed similarly to the VASA-seq data. Here, we kept cells with more than 10^3.5^ and less than 10^6^ uniquely detected transcripts, and with 85%-97% protein-coding transcripts. Cell cycle genes were not removed from the set of highly variable genes, and cell cycle regression was not performed. The effect of the libraries was regressed out, before z-score scaling.

### Comparison between 10X and VASA-seq embryo data

For the comparison, only reads mapping at the 80% 3’ end of gene bodies were used to generate count tables for both VASA and 10x. Only genes expressed in both technologies were used for the comparison. The technology and the number of counts were regressed out from the combined VASA-10x dataset and dimensionality reduction was performed by PCA. Manhattan-based distances between cells were calculated in the combined PCA space. Equivalent clusters were identified as described in the main text. To assign a germ layer to each equivalent cluster, published annotations for the 10x data (Pijuan-Sala et al., 2019) were used (epiblast: epiblast, primitive streak, anterior primitive streak, caudal epiblast, NMP; ectoderm: ExE ectoderm, caudal neurectoderm, rostral neurectoderm, surface ectoderm, forebrain/midbrain/hindbrain, neural crest, spinal cord; mesoderm: nascent mesoderm, caudal mesoderm, ExE mesoderm, intermediate mesoderm, mesenchyme, mixed mesoderm, paraxial mesoderm, pharyngeal mesoderm, somitic mesoderm, cardiomyocytes; endoderm: allantois, def. endoderm, ExE endoderm, gut, parietal endoderm, visceral endoderm; blood: blood progenitors 1, blood progenitors 2, erythroid1, haematoendothelial progenitors, endothelium, erythroid2, erythroid3; PGC: PGC). The prevalent annotation for each equivalent cluster was used.

### Master UMAP for VASA-drop mouse embryo data

The master UMAP, where all cells for all timepoints are integrated together, was obtained as previously described (Wagner et al., 2018). In brief, we first built a directed graph. For each cell in each timepoint, we find the top 30 nearest neighbors in the subset of cells from the same time point and the previous time point (cells from E6.5 are only connected to cells from E6.5). To do so, all the cells in the subset are projected to the PCA space of the latest time-point and distances are calculated using the Manhattan metric. Next, the undirected graph was extracted and used to project the data to the two-dimensional UMAP as previously described (McInnes et al., 2020).

### Expanding the transcriptome annotation

A total of 33,662 demultiplexed and ribo-depleted fastq files for each cell were used to reconstruct the transcriptome and quantify alternative splicing events. To this end, we implemented a custom computational workflow using Snakemake (Köster and Rahmann., 2018) based on Hisat2/StringTie (Pertea et al., 2016) and additional custom scripts. First PCR duplicates were removed through a custom python script that calculates pairwise identity across unique molecular identifiers for each sequenced read within single cells. Then reads were grouped by previously obtained Leiden clusters and mapped to the reference mouse genome assembly, version GRCm38, using HISAT2 (Kim et al., 2019). We performed the alignments implementing the recommended configuration for HISAT2 and genome indexing to ensure an optimal performance during later steps of the transcriptome assembly (Pertea et al., 2016).

The alignments for each cluster were assembled and then merged using StringTie (Pertea et al., 2015). The resulting GTF file is then compared to the input transcriptome annotation using gtfcompare (Pertea and Pertea, 2020) which assigns a classification code to each assembled transcript, which is subsequently used to filter transcripts with codes that indicate additional portions of annotated transcripts or novel genes. Novel transcripts spanning three or more exons that were classified under code k, m, n, j, x, i or y were appended to the input transcriptome annotation, expanding the original set of annotated transcripts. Finally, to further improve the quality of potentially novel transcripts, additional custom filtering steps were implemented to avoid novel transcripts due to false positive novel exons. This filter is particularly important for transcripts assembled from reads that are mapped to repetitive sequences or exons that are ≤30 nt, which can arise from HISAT2 misalignments. To annotate potentially novel microexons, we used MicroExonator, a specialized computational workflow for discovering and quantifying microexons (Parada et al., 2021). After running the MicroExonator’s discovery module we obtained a transcriptome annotation, which was later processed with custom scripts to limit the number of alternative transcription start and end sites.

### Quantification of alternative splicing events across cell-types

The final GTF from the expanded transcriptome annotation was used to quantify isoforms and alternative splicing events using Whippet (Sterne-Weiler et al., 2018). We ran Whippet through MicroExonator’s downstream module to profile alternative splicing events using scRNA-seq data, which enabled randomized aggregations of cells into pseudo-bulks and pairwise comparisons of AS profiles across cell-types. To determine relevant pairwise comparison of AS profiles across cell-types, we used PAGA (Wolf et al., 2019) to calculate connectivities between cell clusters based on gene expression. We then compared the 72 pairs of clusters which have a connectivity >=0.05. For each comparison, cells from each cluster were randomly pooled to form at least three different pseudo-bulks of 200 or fewer cells. To detect reproducible changes of splicing node inclusion across cell-types, random pseudo-bulk pooling and differential inclusion steps were repeated 50 times for each pairwise comparison. As part of MicroExonator’s workflow, the obtained probabilities of each splicing node to be differentially included were used to fit a beta distribution model and calculate CDF-beta values for each event. DISNs were defined as events with CDF-beta values equal or lower than 0.05. To identify SNMs, we calculated the average ψ values for each splicing node across three randomly defined pseudo-bulks samples for each cell-cluster. For splicing nodes where ψ values could be quantified based on at least 10 reads across at least 50 pseudo-bulks, we calculated the Z-score by comparing to all other pseudo-bulks. We considered a splicing node as SNM for a given cell-type if at least two pseudo-bulks had significant Z-scores (p-value less or equal than 0.05) and an absolute difference of at least 0.3 from the mean across all pseudo-bulks. To show some functional consequences of detected AS events for protein function, we used drawProteins package (Brennan, 2018) to draw scaled diagrams of protein domains and other features annotated in UniProt (Apweiler et al., 2004).

## Data and Software Availability

Mapping, analysis scripts and supplementary tables are available at https://github.com/anna-alemany/vasaseq, https://github.com/hemberg-lab/MicroExonator (branch joe), and https://github.com/hemberg-lab/RNA_seq_snakepipes (branch joe).

